# Large-Scale Sparse Regression for Multiple Responses with Applications to UK Biobank

**DOI:** 10.1101/2020.05.30.125252

**Authors:** Junyang Qian, Yosuke Tanigawa, Ruilin Li, Robert Tibshirani, Manuel A. Rivas, Trevor Hastie

## Abstract

In high-dimensional regression problems, often a relatively small subset of the features are relevant for predicting the outcome, and methods that impose sparsity on the solution are popular. When multiple correlated outcomes are available (multitask), reduced rank regression is an effective way to borrow strength and capture latent structures that underlie the data. Our proposal is motivated by the UK Biobank population-based cohort study, where we are faced with large-scale, ultrahigh-dimensional features, and have access to a large number of outcomes (phenotypes): lifestyle measures, biomarkers, and disease outcomes. We are hence led to fit sparse reduced-rank regression models, using computational strategies that allow us to scale to problems of this size. We use an iterative algorithm that alternates between solving the sparse regression problem and solving the reduced rank decomposition. For the sparse regression component, we propose a scalable iterative algorithm based on adaptive screening that leverages the sparsity assumption and enables us to focus on solving much smaller sub-problems. The full solution is reconstructed and tested via an optimality condition to make sure it is a valid solution for the original problem. We further extend the method to cope with practical issues such as the inclusion of confounding variables and imputation of missing values among the phenotypes. Experiments on both synthetic data and the UK Biobank data demonstrate the effectiveness of the method and the algorithm. We present `multiSnpnet` package, available at http://github.com/junyangq/multiSnpnet that works on top of PLINK2 files, which we anticipate to be a valuable tool for generating polygenic risk scores from human genetic studies.

## 1 Introduction

The past two decades have witnessed rapid growth in the amount of data available to us. Many areas such as genomics, neuroscience, economics and Internet services have been producing increasingly larger datasets that have high dimension, large sample size, or both. A variety of statistical methods and computational tools have been developed to accommodate this change so that we are able to extract valuable information and insight from these massive datasets (Hastie et al., 2009; Efron, Hastie, 2016; Dean, Ghemawat, 2008; Zaharia et al., 2010; Abadi et al., 2016).

One major motivating application for this work is the study of data from population-scale cohorts like UK Biobank with genetic data from over one million genetic variants and phenotype data from thousands of phenotypes in over 500,000 individuals (Bycroft et al., 2018). These data present unprecedented opportunities to explore very comprehensive genetic relationships with phenotypes of interest. In particular, the subset of tasks we are interested in is the prediction of a person’s phenotype value, such as disease affection status, based on his or her genetic variants.

Genome-wide association studies (GWAS) is a very powerful and widely used framework for identifying genetic variants that are associated with a given phenotype. See, for example, Visscher et al. (2017) and the references therein. It is based on the results of univariate marginal regression over all candidate variants and tries to find a subset of significant ones. While being computationally efficient and easy to interpret, GWAS has fairly limited prediction performance because at most one predictor can present in the model. If prediction performance is our main concern, it is natural to consider the class of multivariate methods, i.e. that which considers multiple variants simultaneously. In the past, *wide* data were prevalent where only a limited number, like thousands, of samples were available. In this regime, some sophisticated multivariate methods could be applicable, though they have to more or less deal with dimension reduction or variable selection. In this setting, we handle hundreds of thousands samples and even more variables. In such cases, statistical methods and computational algorithms become equally important because only efficient algorithmic design will allow for the application of sophisticated statistical modeling. Recently, we introduced some algorithms addressing these challenges. In particular, Qian et al. (2019) proposed an iterative screening framework that is able to fit the exact lasso/elastic-net solution path in large-scale and ultrahigh-dimensional settings, and demonstrate competitive computational efficiency and superior prediction performance over previous methods.

In this paper, we consider the scenarios where multivariate responses are available in addition to the multiple predictors, and propose a suite of statistical methods and efficient algorithms that allow us to further improve the statistical performance in this large *n* and large *p* regime. Some characteristics we want to leverage and challenges we want to solve include:

### Statistics

There are thousands of phenotypes available in the UK Biobank. Many of them are highly correlated with each other and can have a lot of overlap in their driving factors. By treating them separately, we lose this information that could have been used to stabilize our model estimation. The benefit of building a joint model can be seen from the following simplified model. Suppose all the outcomes **y**^*k*^, *k* = 1, …, *q* are independent noisy observations of a shared factor **u** = **X***β* such that **y**^*k*^ = **u** + **e**^*k*^. It is easy to see that by taking an average across all the outcomes, we obtain a less noisy response 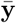, and this will give us more accurate parameter estimation and better prediction than the model built on any of the single outcome. The assumption of such latent structure is an important approach to capturing the correlation structure among the outcomes and can bring in a significant reduction in variance if the data indeed behave in a similar way. We will formalize this belief and build a model on top of it. In addition, in the presence of high-dimensional features, we will follow the “bet on sparsity” principle (Hastie et al., 2009), and assume that only a subset of the predictors are relevant to the prediction.

Therefore, the statistical model we will build features two major assumptions: **low-rank** in the signal and **sparse effect**. Furthermore, we will introduce integrated steps to systematically deal with confounders and missing values.

### Computation

On a large-scale dataset, building a multivariate model can pose great computational challenges. For example, loading the entire UK Biobank dataset into memory with double precision will take more than one terabyte of space, while typically most existing statistical computing tools assume that the data are already sitting in memory. Even if large memory is available, one can always encounter data or construct features so that it becomes insufficient. Hence, instead of expecting sufficient memory space, we would like to find a scalable solution that is less restricted by the size of physical memory.

There is a dynamic data access mechanism provided by the operating system called memory mapping (Bovet, Cesati, 2005) that allows for easy access to larger-than-memory data on the disk. In essence, it carries a chunk of data from disk to memory when needed and swap some old chunks of data out of memory when it is full. In principle, we could add a layer of memory mapping on top of all the procedures and then access the data as if they were in memory. However, there is one important practical component that should never be ignored: disk I/O. This is known to be expensive in the operating system and can greatly delay the computation if frequent disk I/Os are involved. For this reason, we do not pursue first-order gradient-based methods such as stochastic gradient descent (Bottou, 2010) or dual averaging (Xiao, 2010; Duchi et al., 2011) because it can take a large number of passes over the data for the objective function to converge to the optimum.

To address this, we design the algorithm so that it needs as few full passes over the data as possible while solving the exact objective. In particular, by leveraging the sparsity assumption, we propose an adaptive screening approach that allows us to strategically select a small subset of variables into memory, do intensive computation on the subset, and then verify the validity of all the left-out variables. The last step is important because we want to guarantee that the solution obtained from the algorithm is a valid solution to the original full problem.

### 1.1 Reduced-Rank Regression for Multiple Responses

In the standard multivariate linear regression model, given a model matrix **X** = (**x**_1_, …, **x**_*p*_) ∈ ℝ^*n*×*p*^ and a multivariate response matrix **Y** = (**y**_1_, …, **y**_*q*_) ∈ ℝ^*n*×*q*^, we assume that

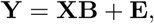

where each row of **E** = (**e**_1_, …, **e**_*q*_) is assumed to be an independent sample from some multivariate Gaussian distribution 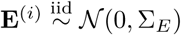. When *n* ≥ *q*, it is easy to see that an maximum likelihood estimator (MLE) can be found by solving a least squares problem with multiple outcomes, i.e.

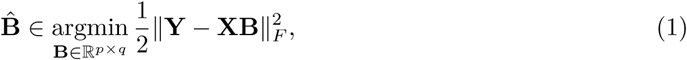

where 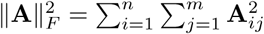 is the squared Frobenius norm of a matrix **A** ∈ ℝ^*n*×*m*^. When *n* ≥ *p* and **X** has full rank, (1) has the closed-form solution 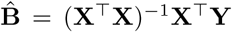. Notice that this is equivalent to solving *q* single-response regression problems separately.

However, in many scenarios, there can be some correlation structure in the signals that we can capture to improve the statistical efficiency of the estimator. One approach to modeling the correlation is to assume that there is a set of latent factors that act as the drivers for all the outcomes. When we assume that the dependencies of the latent factors on the raw features and the outcomes on the latent factors are both linear, it is equivalent to making a low-rank assumption on the coefficient matrix. Reduced-rank regression (Anderson, 1951, hereafter RRR) assumes that the coefficient matrix **B** has a fixed rank *r* ≤ min(*p, q*), or

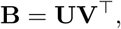

where **U** = (**u**_1_, …, **u**_*r*_) ∈ ℝ^*p*×*r*^, **V** = (**v**_1_, …, **v**_*q*_)^T^ ∈ ℝ^*q*×*r*^.^1^ With the decomposed coefficient matrices, an alternative way to express the multivariate model is to assume that there exists a set of latent factors {**z**_*ℓ*_ ∈ ℝ^*n*^ : 1 ≤ *ℓ* ≤ *r*} such that for each *ℓ*,

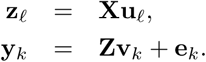

Figure 1 gives a visualization of the dependency structure described above. It can also be seen as a a multilayer perceptron (MLP) with linear activation and one hidden layer, or multitask learning with bottleneck. We notice that under the decomposition, the parameters are not identifiable. In fact, if we apply any nonsingular linear transformation **M** ∈ ℝ^*r*×*r*^ such that **V**^*′*^ = **VM**^T^ and **U**^*′*^ = **UM**^−1^, it yields the same model but different parameters. As a result, we also have an infinite number of MLEs.

**Figure 1:**
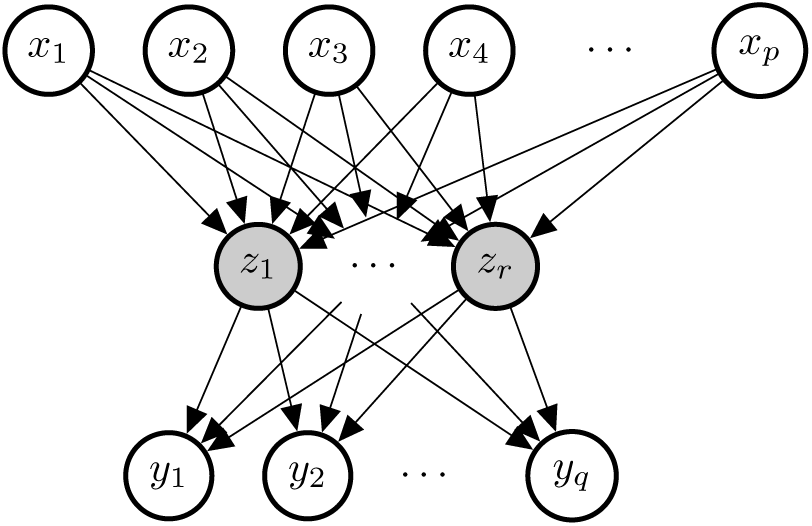
Diagram of the reduced rank regression. The nodes in grey are latent variables. The arrows represent the dependency structure. Known as *multitask* learning in the machine learning community.

Under the rank constraint, an explicit global solution can be obtained. Let **MDN**^T^ be the singular value decomposition (SVD) of 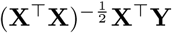, a set of solution is given by **Û** = (**X**^**T**^**X**) ^−1^ **X**^**T**^ **YN**, 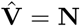. Velu, Reinsel (2013) has a comprehensive discussion on the model under classical large *n* settings.

### 1.2 Sparse Models in High-Dimensional Problems

In the setting of high-dimensional problems where *p* > *n*, the original low-rank coefficient matrix **B** can be unidentifiable. Often sparsity is assumed in the coefficients to model the belief that only a subset of the features are relevant to the outcomes. To find such a sparse estimate of the coefficients, a widely used approach is to add an appropriate non-smooth penalty to the original objective function to encourage the desired sparsity structure. Common choices include the lasso penalty (Tibshirani, 1996), the elastic-net penalty (Zou, Hastie, 2005) or the group lasso penalty (Yuan, Lin, 2006). There has been a great amount of work studying the consistency of estimation and model selection under such settings. See Greenshtein, Ritov (2004); Meinshausen, Bühlmann (2006); Zhao, Yu (2006); Bach (2008); Wainwright (2009); Bickel et al. (2009); Obozinski et al. (2011); Bühlmann, Van De Geer (2011) and references therein. In particular, the group lasso, as the name suggests, encourages group-level sparsity induced by the following penalty term:

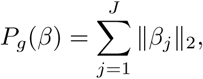

where 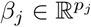 is the subvector corresponding the *j*th group of variables and 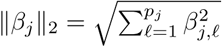 is the vector *ℓ*_2_-norm. The *ℓ*_2_-norm enforces that if the fitted model has 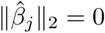, all the elements in 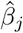 will be 0, and otherwise with probability one all the elements will be nonzero. This yields a desired group-level selection in many applications. Throughout the paper, we will adopt the group lasso penalty, defining each predictor’s coefficients across all outcomes as a distinct group, in order to achieve homogeneous sparsity across multiple outcomes. In addition to variable selection for better prediction and interpretation, we will also see the computational advantages we leverage to develop an efficient algorithm.

## 2 Sparse Reduced-Rank Regression

Given a rank *r*, we are going to solve the following penalized rank-constrained optimization problem:

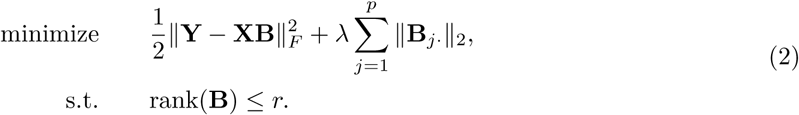

Alternatively, we can decompose the matrix explicitly as **B** = **UV**^T^ where **U** ∈ ℝ^*p*×*r*^, **V** ∈ ℝ^*q*×*r*^. It can be shown that the problem above is equivalent to the Sparse Reduced Rank Regression (SRRR) proposed by Chen, Huang (2012):

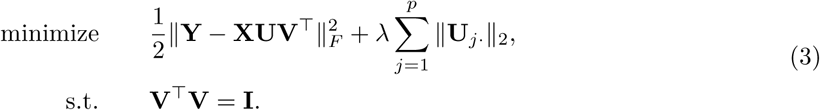

Alternating minimization was proposed by Chen, Huang (2012) to solve this non-convex optimization problem, where two algorithms were considered: subgradient descent and a variational method.

The subgradient method was shown to be faster when *p* ≫ *n* and the variational method faster when *n* ≫ *p*. However, in each iteration, the computational complexity of either method is at least quadratic in the number of variables *p*. It makes the problem almost intractable in ultrahighdimensional problems, which is common, for example, in modern genetic studies. Moreover, to obtain a model with good prediction performance, we are interested in solving the problem over multiple *λ*’s rather than a single one. For such purposes, we design a path algorithm with adaptive variable screening that will be both memory and computationally efficient.

## 3 Fast Algorithms for Large-Scale and Ultrahigh-Dimensional Problems

First, we present a naive version of the path solution, which will be the basis of our subsequent development. The path is defined on a decreasing sequence of *λ* values *λ*_max_ = *λ*_1_ > *λ*_2_ >… > *λ*_*L*_ ≥ 0, where *λ*_max_ is often defined by one that leads to the trivial (e.g. all zero) solution and the rest are often determined by an equally spaced array on the log scale. In particular, for Problem (2), we are able to figure out the exact lower bound of *λ*_max_ for which the solution is trivial.

### Lemma 1.

*In problem (2), if r* > 0, *the maximum λ that results in a nontrivial solution* 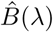 *is*

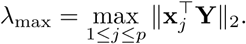

The proof is straightforward, which is a result of the Karush–Kuhn–Tucker (KKT) condition (See Boyd et al. (2004) for more details). We present the full argument in Appendix A.1. The naive path algorithm tries to solve the problem independently across different *λ* values.

### 3.1 Alternating Minimization

The algorithm is described in Algorithm 1. For each *λ* value, it applies alternating minimization to Problem (3) till convergence.

In the V-step (4), we will be solving the orthogonal Procrustes problem given a fixed **U**^(*k*)^. An explicit solution can be constructed from the singular value decomposition, as detailed in the following Lemma.

#### Lemma 2.

*Suppose p* ≥ *r and* **Z** ∈ ℝ^*p*×*r*^. *Let* **Z** = **MDN**^T^ *be its (skinny) singular value decomposition, where* **M** ∈ ℝ^*p*×*r*^, **D** = ℝ^*r*×*r*^ *and* **N** ∈ ℝ^*r*×*r*^. *An optimal solution to*

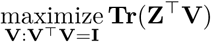

*is given by* 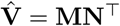, *and the objective function has optimal value ‖***Z‖** _*\**_, *the nuclear norm of* **Z**.

*Proof*. See in Appendix A.2. □

To analyze the computational complexity of the algorithm, we see a one-time computation of **Y**^T^**X** that costs *O*(*npq*). In each iteration, there is *O*(*pqr*) complexity for the matrix multiplication **Y**^T^**XU**^(*k*)^ and *O*(*qr*^2^) for computing the SVD and the final solution. Therefore, the per-iteration computational complexity for the V-step is *O*(*pqr* + *qr*^2^), or *O*(*pqr*) when *p* ≫ *q*.

#### Algorithm 1 Alternating Minimization

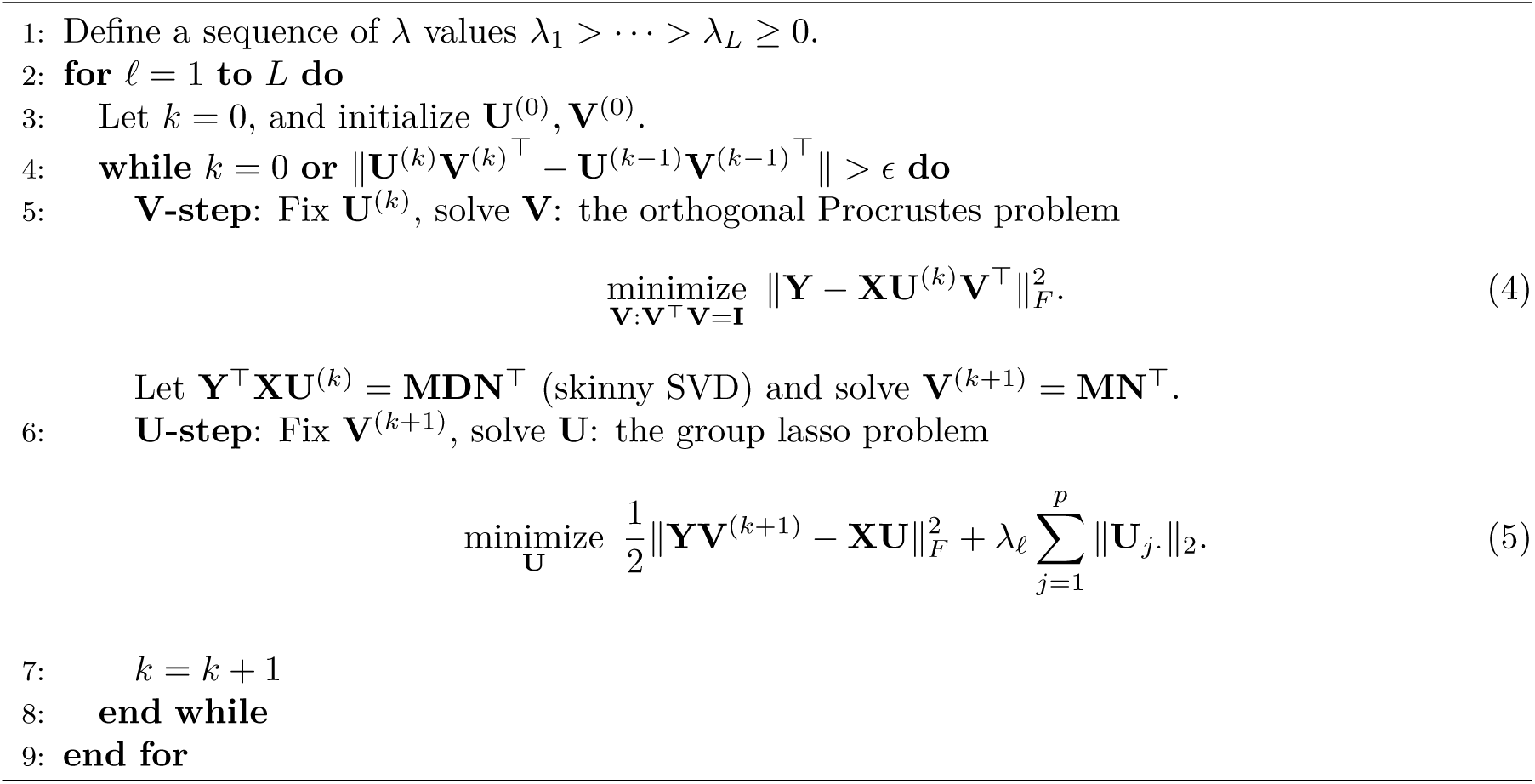

In the U-step, we are solving a group lasso problem. Computing **YV**^(*k*+1)^ takes *O*(*nqr*) time. The group-lasso problem can be solved by **glmnet** (Friedman et al., 2010) with the `mgaussian` family. With coordinate descent, its complexity is 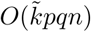, where 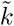 is the number of iterations until convergence and is expected to be small with a reasonable initialization, for example, provided by warm start. Thus, the per-iteration complexity for the U-step is 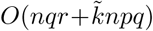, which is 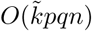 when *p* ≫ *r*.

Therefore, the overall computational complexity scales at least linearly with the number of features, and will have a large multiplier if the sample size is large as well. While subsampling can effectively reduce the computational cost, in high-dimensional settings, it is critical to have sufficient samples for the quality of estimation. Instead, we seek for computational techniques that can lower the actual number of features involved in expensive iterative computation without giving up any statistical efficiency. Thanks to the induced sparsity by the objective function, we are able to achieve it by variable screening.

### 3.2 Variable Screening for Ultrahigh-Dimensional Problems

In this section, we discuss strategic ways to find a good subset of variables to focus on in the computation that would allow us to reconstruct the full solution easily. In particular, we would like to iterate through the following steps for each *λ*:

1. **Screen** a strong set *S* and treat all the left-out variables *S*^*c*^ as null variables that potentially have zero coefficients;
2. **Solve** a significantly smaller problem on the subset of variables *S*;
3. **Check** an optimality condition to guarantee the constructed full solution 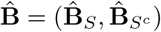 with 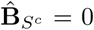 is indeed a valid solution to the original problem. If the condition is not satisfied, go back to the first step with an expanded set *S*.

#### 3.2.1 Screening Strategies

We have seen Lemma 1 that determines the entry point of any nonzero coefficient on the solution path. Furthermore, there is evidence that the variables entering the model (as one decreases the *λ* value) tend to have large values by this criterion. Tibshirani et al. (2012) developed on this idea and proposed the strong rules as a sequential variable screening mechanism. The strong rules state that in a standard lasso problem with the model matrix **X** = (**x**_1_, …, **x** _*p*_) ∈ ℝ^*n*×*p*^ and a single response **y** ∈ ℝ^*n*^, assume 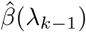 is the lasso solution at *λ*_*k*−1_, then the *j*th predictor is discarded at *λ*_*k*_ if

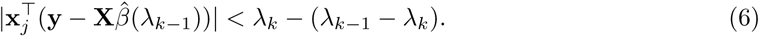

The key idea is that the inner product above is almost “non-expansive” in terms of *λ*. As a result, the KKT condition suggests that the variables to be discarded by (6) would have coefficient 0 at *λ*_*k*_. However it is not a guarantee. The strong rules can fail, though failures occur rarely when *p* > *n*. In any case, the KKT condition is checked to ensure the exact solution is found. Although Tibshirani et al. (2012) focused mostly on the lasso-type problem, they also suggested extension to general objective functions and penalties. For general objective function *f* (*β*) with *p*_*j*_-norm penalty 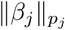 for the *j*th group, the screening criterion will be based on the dual norm of its gradient 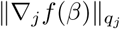 where 1*/p*_*j*_ + 1*/q*_*j*_ = 1.

Inspired by the general strong rules, we propose three sequential screening strategies for the sparse reduced rank objective (3), named after their respective characteristics: Multi-Gaussian, Rank-Less and Fix-V. They are based either on the solution of a relaxed convex problem at the same *λ*_*k*_ or on the exact solution at the previous *λ*_*k*−1_.

- (Multi-Gaussian) Solve the full-rank convex problem at *λ*_*k*_ and use its active set as the candidates for the low-rank settings. The main advantage is that the screening is always stable due to the convexity. However this approach often overselects and brings extra burden to the computation. By assuming a higher rank than necessary, the effective number of responses would become more than that of a low-rank model. As a result, more variables would potentially be needed to serve for an enlarged set of responses.
- (Rank-Less) Find variables that have large 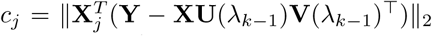. This is analogous to the strong rules applied to the vanilla multi-response lasso ignoring the rank constraint.
- (Fix-V) Find variables that have large 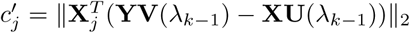. This is similar to the strong rules applied in the **U**-step with **V** assumed fixed. To see the rationale better, we take another perspective. The squared error in SRRR (3) can also be written as

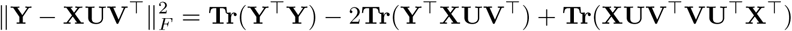

Since **V**^T^**V** = **I**, the optimization problem becomes

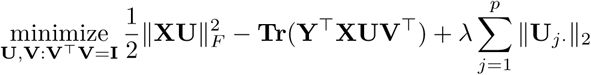

For any given **U**, we can solve **V** = **MN**^T^, where **Y**^T^**XB** = **MDN**^T^ is its singular value decomposition. Let 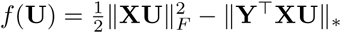. The problem is reduced to

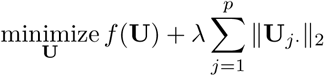

The general strong rule tells us to screen based on the gradient; that is

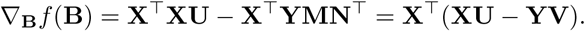

Therefore, the general strong rules endorse the use of this screening rule.

We do some experiments to compare the effectiveness of the rules. We simulate the model matrix under an independent design and an equi-correlated design with correlation *ρ* = 0.5. The true solution path is computed using Algorithm 1 with several random initializations and the convex relaxation-based initialization (as in the Multi-Gaussian rule). Let *S*(*λ*) be the true active set at *λ*. For each method *ℓ* above, we can find, based on either the exact solution at *λ*_*k*_−_1_ or the fullrank solution at *λ*_*k*_, the threshold it needs so that by the screening criterion, the selected subset *Ŝ*(*λ*_*k*_)^(*ℓ*)^ contains the true subset at *λ*_*k*_, i.e. *Ŝ*(*λ*_*k*_|*λ*_*k*−1_)^(*ℓ*)^ *⊇ S*(*λ*_*k*_). This demonstrates how deep each method has to search down the variable list to include all necessary variables, and thus how accurate the screening mechanism is — the larger the subset size, the worse the method is.

We see from both plots that the curve of the Fix-V method is able to track that of the exact subset fairly well, while the Rank-Less and Multi-Gaussian methods both choose a much larger subset in order to cover the subset of active variables in the exact solution. In the rest of the paper, we will adopt the Fix-V method to do variable screening.

#### 3.2.2 Optimality Condition

Although the Fix-V method turns out to be most effective in choosing the subset of variables, in practice we have no access to the true subset and have to take an estimate. Instead of trying to find a sophisticated threshold, we will do batch screening at a fixed size (this size can change adaptively though). Given a size *K*, we will take the *K* variables that rank the top under this criterion. Clearly we can make mistakes by having left out some important variables in the screening stage. In order to make sure that our solution is exact rather than approximate in terms of the original problem, we need to check the optimality condition and take in more variables when necessary.

Suppose we find a solution **Û**_*S*_, 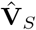 on a subset of variables **X**_*S*_ by alternating minimization. We will verify the assembled solution 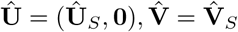 is a limit point of the original optimization problem. The argument is supported by the following lemma.

##### Lemma 3.

*In the U-step (12), given* **V** *and λ, if we have an exact solution* **Û**_*S*_ *for the sub-problem with* **X**_*S*_, *then* **Û** = (**Û**_*S*_, **0**) *is a solution to the full problem if and only if for all j* ∈ *S*^*c*^,

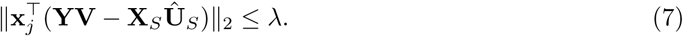

*Proof*. Since this is a convex problem, **Û** is a solution if and only if **0** ∈ *∂f* (**Û**) where *f* is the objective function in (12) and *∂f* is its subdifferential. For the vector *ℓ*_2_-norm, we know that the subdifferential of ‖**x**‖ _2_ is {**s** ∈ ℝ^*p*^ : ‖**s**‖ _2_ ≤ 1} if **x** = **0** and {**x**/‖**x**‖_2_} if **x** ≠ **0**. Notice that **X**_*S*_ **Û**_*S*_ = **XÛ** by the definition of **Û**. Since we have an exact solution on *S*, we know **0** ∈ *∂f* (**Û**)_*j·*_ for all *j* ∈ *S*. On the other hand, for *j* ∈ *S*^*c*^, 0 ∈ *∂f* (**Û**) if and only if 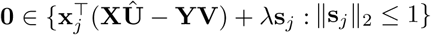, which is further equivalent to 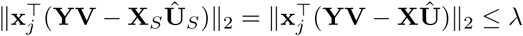 □

Therefore, once we obtain a solution **Û**_*S*_, 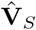 for the sub-problem and get condition (7) verified, we know in the V-step, by the lemma above, **Û** = (**Û**_*S*_, **0**) is the solution given 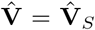. In the U-step, since **XÛ** = **X**_*S*_ **Û**_*S*_, **Û** is the solution to the full problem. We see that (**Û**, 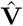) is a limiting point of the alternating minimization algorithm for the original problem. However if the condition fails, we expand the screened set or bring in the violated variables, and do the fit again. We should note that when we say an exact solution to the original problem, we do not claim it to be a local minimum or global minimum, unless under some regularity conditions as will be briefly discussed later. It is a limiting point of the vanilla alternating minimization algorithm, i.e. Algorithm 1. In other words, if we start from the constructed solution (with zero coefficients for the leftout variables), the algorithm should converge in one iteration and return the same solution.

We have seen the main ingredients of the iterative algorithm: screening, solving and checking. Next we discuss some useful practical considerations and extensions.

### 3.3 Computational Considerations

#### 3.3.1 Initialization and Warm Start

Recall that in the training stage our goal is to fit an SRRR solution path across different *λ* values. It is easy to see that with a careful choice of parameterization, the path is continuous in *λ*. To leverage this property, we adopt a warm start strategy. Specifically, we initialize the coefficients of the existing variables at *λ*_*k*+1_ using the solution at *λ*_*k*_ and zero-initialize the newly added variables. With warm start, much less iterations will be needed to converge to the new minimum.

However, this by no means guarantees that we are all on a good path. It’s likely that we are trapped into a neighborhood of local optimum and end up with much higher function value than the global minimum. One way to alleviate this, if affordable, is to solve the corresponding full-rank problem first, and initialize the coefficients with low-rank approximation of the full-rank solution. We can compare the limiting function values with the warm-start initialization and see which converges to a better point. Although we didn’t use in the actual implementation and experiments, one could also do random exploration — randomly initialize some of the coefficients, run the algorithm multiple times and find one that achieves the lowest function value. That said, we lose the advantage of warm start though. The good news is, in the experiments we have done, we didn’t observe very clear suboptimal behavior by the warm start and full-rank strategies.

#### 3.3.2 Early Stopping

Although we pre-specify a sequence of *λ* values *λ*_1_ > *λ*_2_ > …> *λ*_*L*_ where we want to fit the SRRR models, we do not have to fit them all given our goal is to find the best predictive model. Once the model starts to overfit as we move down the *λ* list, we can stop our process since the later models will have no practical use and are expensive to train. Therefore, in the actual computation, we monitor the validation error along the solution path and call it a stop if it shows a clear upward trend. One other point we would like to make in this regard is that the validation metric can be defined either as an average MSE over all phenotypes or a subset of phenotypes we are most interested in. This is because practically the best *λ* value can be different for different phenotypes in the joint model.

### 3.4 Extensions

#### 3.4.1 Standardization

We often want to standardize the predictors if they are not on the same scale because the penalty term is not invariant to change of units of the variables. However we emphasize that some thought has to be put into this before standardizing the predictors. If the predictors are already on the same scale, standardizing them could bring unintended advantages to variables with smaller variance themselves. It is more reasonable not to standardize in such cases.

In terms of the outcomes, since they can be at different scales, it is important to standardize them in the training stage so that no one dominates in the objective function. At prediction (both training and test time), we scale back to the original levels using their respective variances from the training set. In fact, the real impact an outcome has to the overall objective is determined by the proportion of unexplained variance. It would be good to weight the responses properly based on this if such information is available or can be estimated, e.g. via heritability estimation for phenotypes in genetic studies.

#### 3.4.2 Weighting

Sometimes we have strong reasons or evidence to prioritize some of the predictors than the others. We can easily extend the standard objective (3) and reflect this belief in a weighted penalty 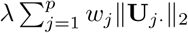 where the weight *w*_*j*_ controls inversely the relative importance of the *j*th variable. For example, *w*_*j*_ = 0 implies *j*th variable will always be included in the model, while a large *w*_*j*_ will almost exclude the variable from the model.

In the response space, we can also impose a weighting mechanism to priortize the training of certain responses. For a given set of nonnegative weights *w*_*k*_, 1 ≤ *k* ≤ *q*, the SRRR objective (3) can be modified to 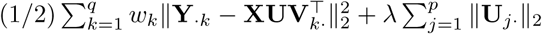 with the same constraint, or equivalently,

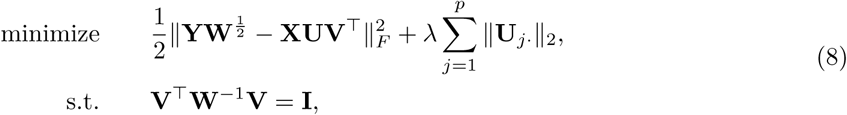

where the weight matrix **W** = **diag**(*w*_1_, …, *w*_*q*_). To solve the problem with our alternating minimization scheme, we can see that in the V-step, instead of solving the standard orthogonal Procrustes problem with an elegant analytic solution derived from the SVD, we have to deal with a so-called weighted orthogonal Procrustes problem (WOPP). Finding the solution of the WOPP is far more complicated. See, for instance, Mooijaart, Commandeur (1990), Chu, Trendafilov (1998) and Viklands (2006). An iterative procedure is often needed to compute the solution. For better computational efficiency, we instead solve the problem with the original orthonormal constraint:

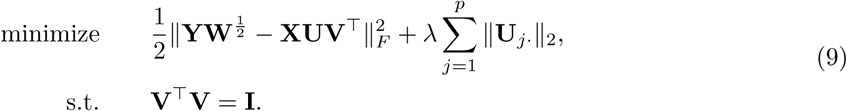

That is, we amplify the magnitude of some responses so that the objective value is more sensitive to the loss incurred on these responses. When making prediction, we will need to scale them back to the original units.

#### 3.4.3 Adjustment Covariates

In some applications such as genome-wide association studies (GWAS), there may be confounding variables **Z** ∈ ℝ^*n*×*m*^ that we want to adjust for in the model. For example, population stratification, defined as the existence of a systematic ancestry difference in the sample data, is one of the common factors in GWAS that can lead to spurious discoveries. This can be controlled for by including some leading principal components of the SNP matrix as variables in the regression (Price et al., 2006). In the presence of such variables, we solve the following problem instead. With a slight abuse of notation, in this section, we use **W** to denote the coefficient matrix for the covariates instead of a weight matrix:

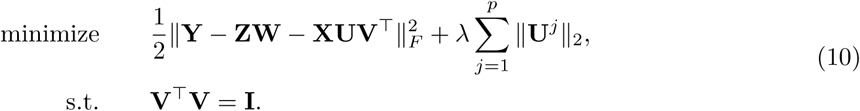

The main components don’t change except two adjustments. When determining the starting *λ* value, we use Lemma 4.

##### Lemma 4.

*In problem (10), if r* > 0, *the maximum λ that results in a nontrivial solution* 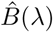 *is*

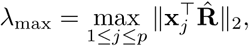

*where* 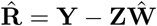 *and* **Ŵ** *is the multiple outcome regression coefficient matrix*.

The proof is almost the same as before. The other nuance we should be careful about is when fitting the model, we should leave those covariates unpenalized because they serve for the adjustment purpose and should not be experiencing the selection stage. In particular, in the U-step (group lasso) given **V**, direct computation would reduce to solving the problem

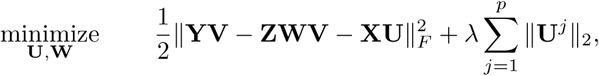

which is not as convenient as standard group lasso problem. Instead, we find that **W** can always be solved explicitly in terms of other variables. In fact, the minimizer **Ŵ** = (**Z**^T^**Z**)^−1^**Z**^T^(**Y** − **XUV**^T^). Plug in and we find that the problem to be solved can be written as

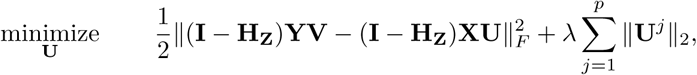

where **H**_**Z**_ = **Z**(**Z**^T^**Z**)^−1^**Z**^T^ is the projection matrix on the column space of **Z**. This becomes a standard group lasso problem and can be solved by using, for example, the **glmnet** package with the mgaussian family.

#### 3.4.4 Missing Values

In practice, there can be missing values in either the predictor matrix or the outcome matrix. If we only discard samples that have any missing value, we could lose a lot of information. For the predictor matrix, we could do imputation as simple as mean imputation or something sophisticated by leveraging the correlation structure. For missingness in the outcome, there is a natural way to integrate an imputation step seamlessly with the current procedure, analogous to the softImpute idea in Mazumder et al. (2010). We first define a projection operator for a subset of two dimensional indices Ω *⊆* {1, …, *n*} × {1, …, *p*}. Let *𝒫*_Ω_ : ℝ^*n*×*p*^ →ℝ^*n*×*p*^ be such that

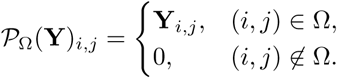

Let Ω be the set of indices where the response values are observed; in other words, Ω^*c*^ is the set of missing locations. Instead of (3), now we solve the following problem.

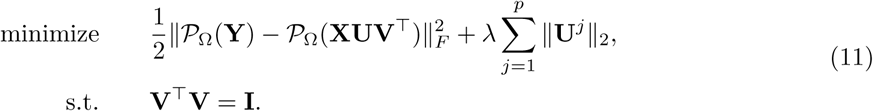

We can easily see that an equivalent formulation of the problem is

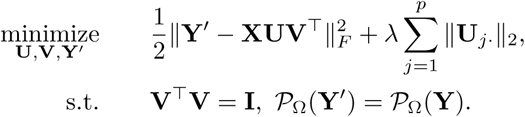

This inspires a natural projection step to deal with the additional constraint. It can be well integrated with the current alternating minimization scheme. In fact, after each alternation between the U-step and the V-step, we can impute the missing values from the current predictions **XUV**^T^, and then continue into the next U-V alternation with the completed matrix.

#### 3.4.5 Lazy Reduced Rank Regression

There is an alternative way to find a low-rank coefficient profile for the multivariate regression. Instead of pursing to solve the non-convex problem (3) directly, we can follow a two-stage procedure:

1. Solve a full-rank multi-gaussian sparse regression, i.e.,

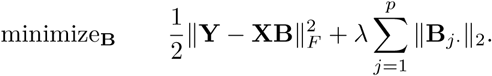
2. Conduct SVD of the resulting coefficient matrix 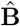 and use its rank *r* approximation as our final estimator.

The advantage of this approach is that it is stable. The first stage is a convex problem and can be handled efficiently by, for example, `glmnet`. A variety of adaptive screening rules are also applicable in this situation to assist dimension reduction. The second stage is fairly standard and efficient as long as there are not too many active variables. However, the disadvantage is clear too. The low-rank approximation is conducted in an unsupervised manner, so could lead to some degrade in the prediction performance.

That said, as before, we should still evaluate the out-of-sample performance as the penalty parameter *λ* varies and pick the best on the solution path as our final estimated model. In many cases, we compute the full-rank model under the exact mode anyways, so the set of lazy models can be thought of as an efficient byproduct for our choice.

### 3.5 Full Algorithm

We incorporate the options above and present the full algorithm in Algorithm 2.

#### Algorithm 2 Large-scale and Ultrahigh-dimensional Sparse Reduced Rank Regression

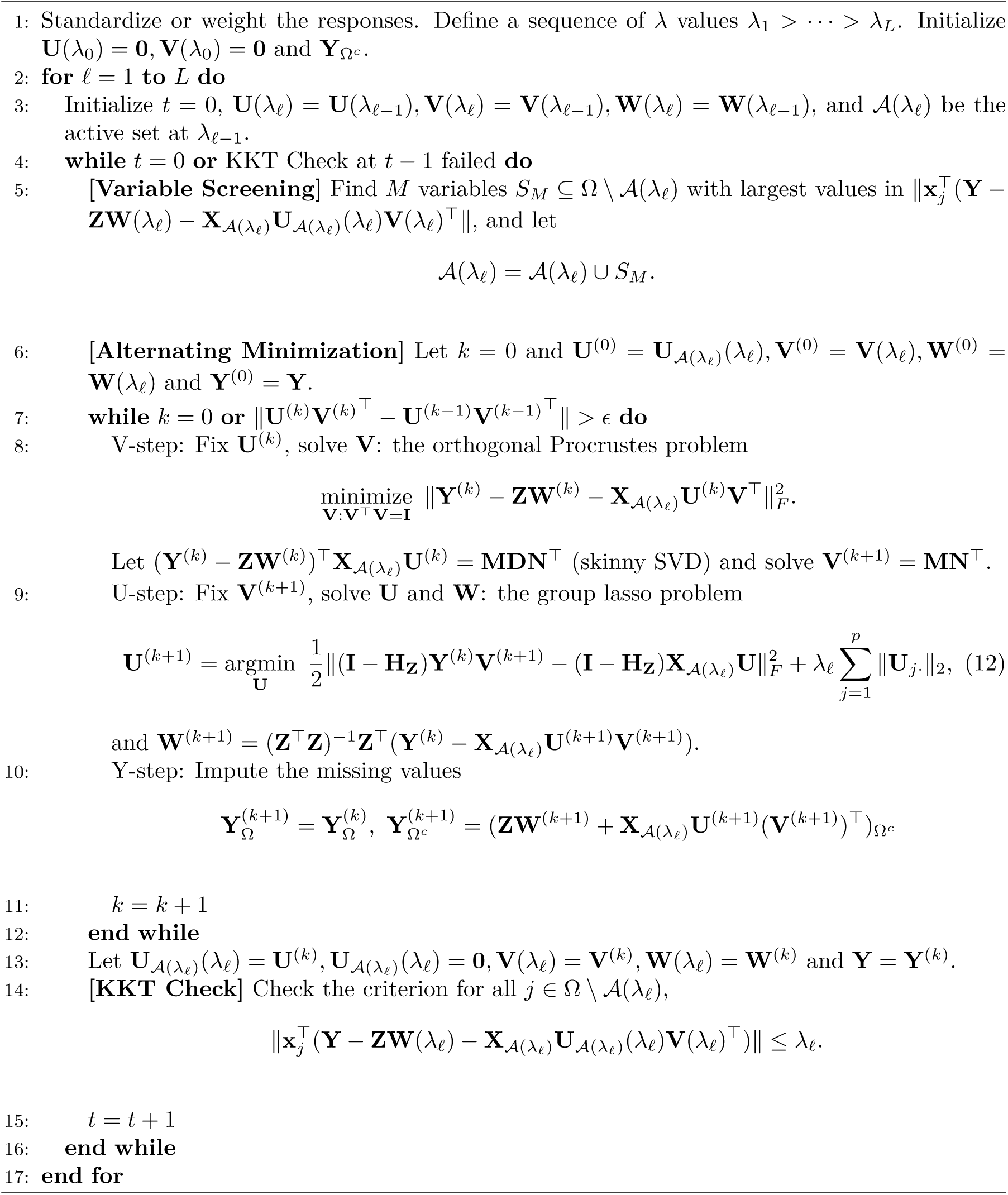

## 4 Convergence Analysis

In this section, we present some convergence properties of the alternating minimization algorithm (Algorithm 1) on sparse reduced rank regression. Let

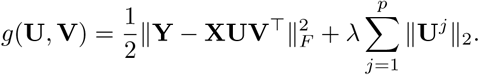

### Theorem 1.

*For any k* ≥ 1, *the function values are monotonically decreasing:*

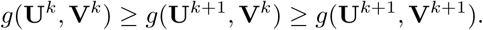

*Furthermore, we have the following finite convergence rate:*

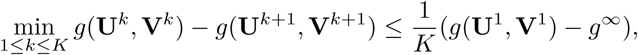

*where g*^*∞*^ = lim_*k*→*∞*_ *g*(**U**^*k*^, **V**^*k*^). *It implies that the iteration will terminate in O*(1*/ϵ*) *iterations*.

The proof is straightforward and we won’t detail here. It presents the fact that alternating minimization is a descent algorithm. In fact, this property holds for all alternating minimization or more general blockwise coordinate descent algorithms. However it does not say how good the limiting point is. In the next result, we show a local convergence result that under some regularity conditions, if the initialization is closer enough to a global minimum, it will converge to a global minimum at linear rate. It is based on similar results on proximal gradient descent by Dubois et al. (2019). To define a local neighborhood, it would be easier if we eliminate **V** by always setting it to a minimizer given **U**. That is, the objective function becomes 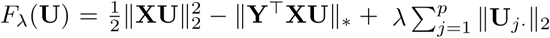. We define a sublevel set *𝒮*_*c*_(*λ*) = {**U** ∈ ℝ^*p*×*r*^ : *F*_*λ*_(**U**) ≤ *c*}.

### Theorem 2.

*Assume* **X**^T^**X** *is invertible and* 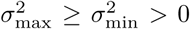 *be its smallest and largest eigenvalues. Let s*_*j*_ *be the jth singular value of* 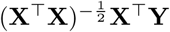. *There exists* 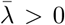 *such that for all* 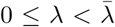 *and* 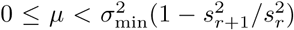, *there is a sublevel set S*(*λ, µ*) *where the level depends on λ and µ such that if* **U**^*k*^ ∈ *𝒮*(*λ, µ*), *we have*

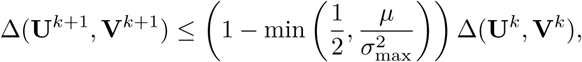

*where* Δ(**U, V**) = *g*(**U, V**) − *g*(**U**^*\**^, **V**^*\**^) *and* (**U**^*\**^, **V**^*\**^) *is a global minimum*.

From a high level, the proof is based on the fact that under the conditions, the function is strongly convex near the global minima. If we starting from this region, we achieve good convergence rate with alternating minimization algorithm. The full proof is given in Appendix A.3.

It is easy to see that the theorem above implicitly assumes the classical setting where *n* ≥ *p* since otherwise **X**^T^**X** would not be invertible. However, it is still applicable to our algorithm. The algorithm does not attempt to solve alternating minimization at the full scale, but only does it after variable screening. With screening, it is very likely that we will again be working under the classical setting. Moreover, with warm start, there is higher chance that the initialization lies in the local region as defined above. Therefore, this theorem can provide useful guidance on the practical computational performance of the algorithm.

## 5 Simulation Studies

We conduct some experiments to gain more insight into the method and compare with the single-response lasso method. Due to space limit, we demonstrate the results in one experiment setting and include results for other settings such as correlated features, deviation from the true low-rank structure etc., in Appendix C. We experiment with three different sizes and three different signal-to-noise ratio (SNR): (*n, p, k*) = (200, 100, 20), (200, 500, 20), (200, 500, 50), where *k* is the number of variables with true nonzero coefficients, and the target SNR = 0.5, 1, or 3. The number of responses *q* = 20 and the true rank *r* = 3. We generate the **X** ∈ ℝ^*n*×*p*^ with independent samples from some multivariate Gaussian *𝒩*(0, Σ_*X*_) where Σ_*X*_ = **I**_*p*_ in this section. More results under correlated designs are presented in the appendix. The response is generated from the true model **Y** = **XUV**^T^ + **E**, where each entry in the support of **U** ∈ ℝ^*p*×*r*^ (sparsity *k*) is independently drawn from a standard Gaussian distribution, and **V** ∈ ℝ^*q*×*r*^ takes the left singular matrix of a Gaussian ensemble. Hence **B** = **UV**^T^ is the true coefficient matrix. The noise matrix is generated from 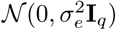, where 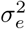 is chosen such that the signal-to-noise ratio

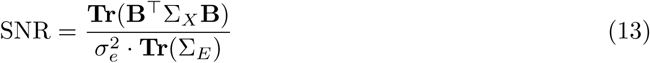

is set to a given level. The performance is evaluated by the test *R*^2^, defined as follows:

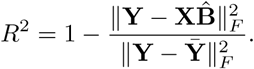

The main insight we obtain from the experiments is that the method is more robust to over-estimating than underestimating the rank. A significant degrade in performance can be identified even if we are only off the rank by 1 from below. In contrast, the additional variance brought along by overestimating the rank doesn’t seem to be a big concern. This, in essence, can be ascribed to bias and variance decomposition. In our settings, the bias incurred in underestimating the rank and thus 1/3 loss of parameters contributes a lot more to the MSE compared with the increased variance due to 1/3 redundancy in the parameters.

## 6 Real Data Application: UK Biobank

The UK Biobank (Bycroft et al., 2018) is a large, prospective population-based cohort study with individuals collected from multiple sites across the United Kingdom. It contains extensive genetic and phenotypic detail such as genome-wide genotyping, questionnaires and physical measures for a wide range of health-related outcomes for over 500,000 participants, who were aged 40-69 years when recruited in 2006-2010. In this study, we are interested in the relationship between an individual’s genotype and his/her phenotypic outcomes. While genome-wide association studies (GWAS) focus on identifying SNPs that may be marginally associated with the outcome using univariate tests, we would like to leverage the additive effect of all SNPs to make good prediction. Recently there is a line of work (Qian et al., 2019; Sinnott-Armstrong et al., 2019; Lello et al., 2018) that builds a lasso solution on the large dataset and shows that the prediction is much improved over previous methods. Furthermore, a number of phenotypes present nontrivial correlation structures and we would like to further improve the prediction and stabilize the variable selection by building a joint model for multiple outcomes.

We focused on 337,199 White British unrelated individuals out of the full set of over 500,000 from the UK Biobank dataset (Bycroft et al., 2018) that satisfy the same set of population stratification criteria as in DeBoever et al. (2018). Each individual has up to 805,426 measured variants, and each variant is encoded by one of the four levels where 0 corresponds to homozygous major alleles, 1 to heterozygous alleles, 2 to homozygous minor alleles and NA to a missing genotype. In addition, we have available covariates such as age, sex, and forty pre-computed principal components of the SNP matrix. Among them, we use age, sex and the top 10 PCs for the adjustment of population stratification (Price et al., 2006).

There are binary responses in the data such as many disease outcomes. Although in principle we can solve for a mixture of Gaussian and binomial likelihood using Newton’s method, for ease of computation in this large-scale setting, it is a reasonable approximation to treat them as continuous responses and fit the standard SRRR model. However, after the model is fit, we will refit a logistic regression on the predicted score to obtain a probability estimation. Notice that the refit is still trained on the training set at each *λ* value.

The number of samples is large in the UK Biobank dataset, so we afford to set aside an independent validation set without resorting to costly cross-validation to find an optimal regularization parameter. We also leave out a subset of observations as test set to evaluate the final model. In particular, we randomly partition the original dataset so that 70% is used for training, 10% for validation and 20% for test. The solution path is fit on the training set, whereas the desired regularization is selected on the validation set, and the final model is evaluated on the test set.

In the experiment, we compare the performance of the multivariate-response SRRR model with the single-response lasso model. To fit the lasso model, we rely on fast implementation of the **snpnet** package (Qian et al., 2019), and we also refer to the lasso results as snpnet in the results section. For continuous responses, we evaluate the prediction by R-squared (*R*^2^). Given a linear coefficient vector 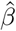 (fitted on the training set) and a subset of data {(*x*_*i*_, *y*_*i*_), 1 ≤ *i* ≤ *n*}, it is defined as

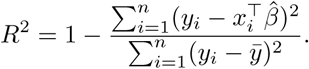

We compute *R*^2^ respectively on the training, validation and test sets. For binary response, misclassification error could be used but it would depend on the calibration. Instead the receiver operating characteristic (ROC) curve provides more information and demonstrates the tradeoff between true positive and false positive rates under different thresholds. The area under the curve (AUC) computes the area under the ROC curve — a larger value indicates a generally better classifier. Therefore, we will evaluate AUCs on the training, validation and test sets for binary responses. When comparing different methods, we evaluate both absolute change and relative change over the baseline method (in particular the already competitive lasso in our case), where the relative change for a given metric is defined as (metric_new_ − metric_lasso_)*/* |metric_lasso_|.

Computationally, in the UK Biobank experiments, the SNP data are stored in a compressed PLINK format with two-bit encodings. PLINK 2.0 (Chang et al., 2015) provides an extensive set of efficient operations including very fast, multithreaded matrix multiplication. In particular, this matrix multiplication module is heavily used in the steps of screening and KKT check in this work and other lasso-based results (Li et al., 2020; Qian et al., 2019) on the UK Biobank.

### 6.1 Asthma and 7 Blood Biomarkers

Here, we defined asthma based on a mixture of self-reported questionnaire data and hospital inpatient record data described in DeBoever et al. (2018); Tanigawa et al. (2019). Furthermore, we focused on 7 additional blood count measurements from Category 100081 in UK Biobank containing results of haematological assays that were performed on whole blood.

We apply the SRRR to the set of phenotypes and expect some performance improvement by leveraging the correlation structure. Choice of the phenotypes: monocyte count, neutrophill count, eosinophill count, basophill count, forced vital capacity (FVC), peak expiratory flow (PEF), and forced expiratory volume in 1 second (FEV1).

Overall, we see small rank representation can maintain predictive power for specific phenotypes (see Figure 4) and that overall the multiresponse model improves the prediction over the single-response lasso model (see Figure 5).

**Figure 2:**
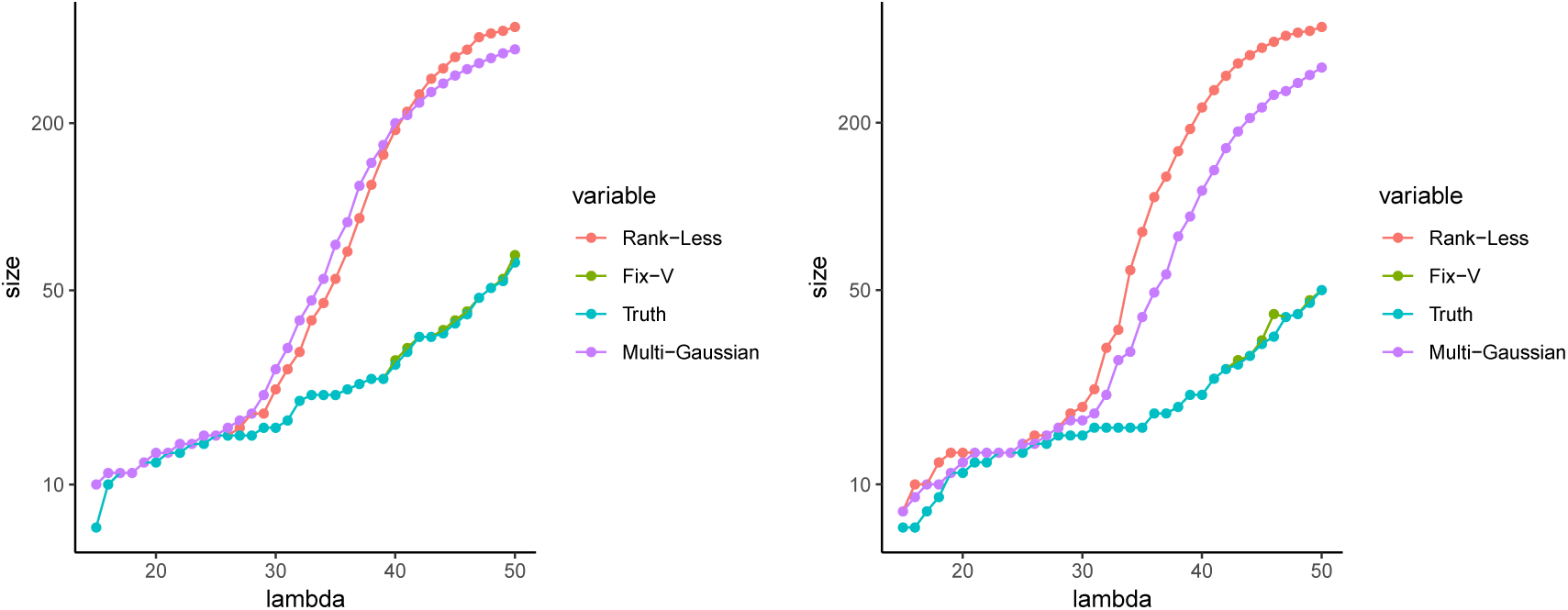
Size of screened set under different strategies. Left: independent design. Right: equi-correlated design with *ρ* = 0.5. Signal-to-noise ratio (SNR) = 1, and we use the true rank = 3.

**Figure 3:**
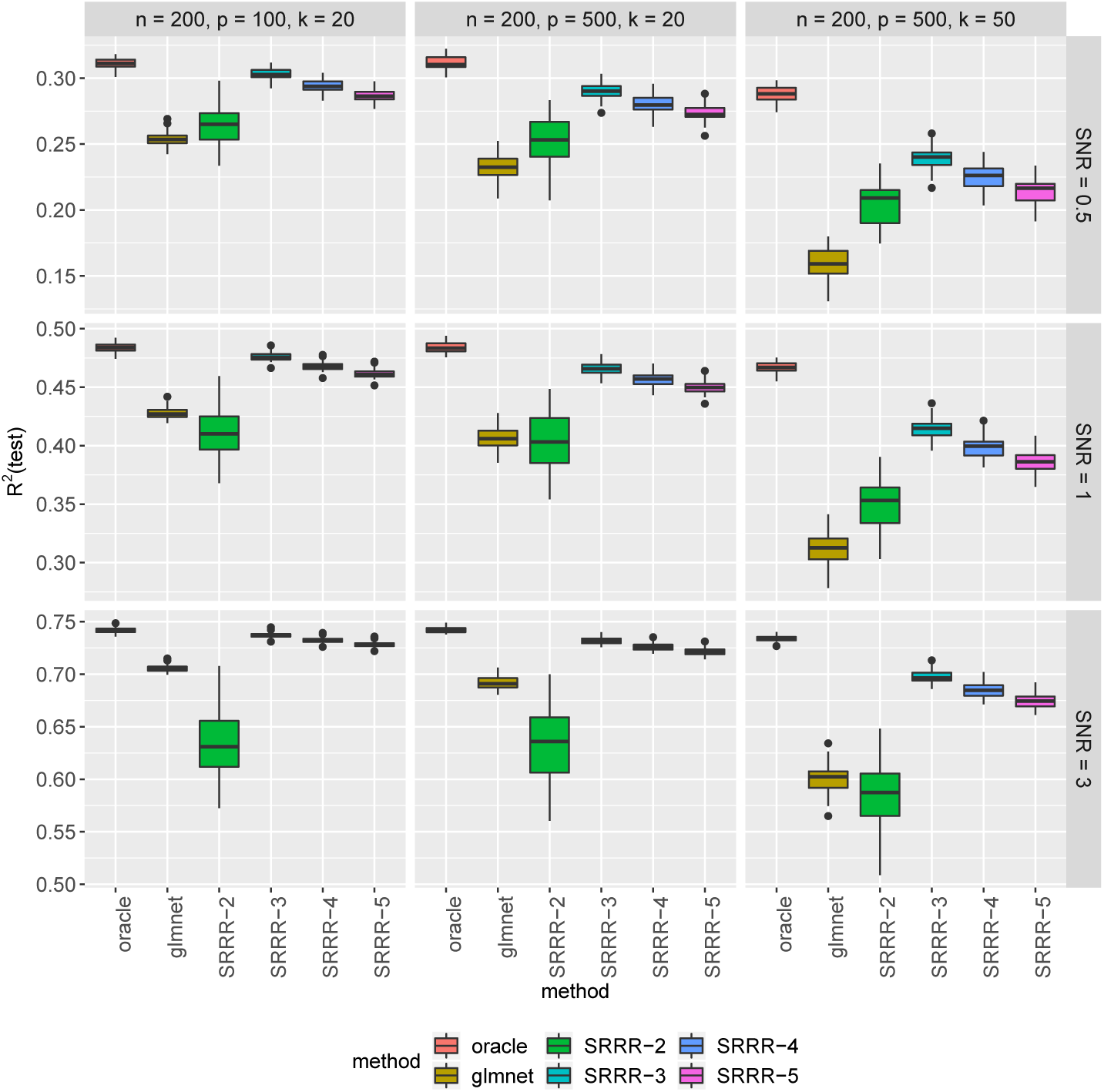
*R*^2^ each run is evaluated on a test set of size 5000. “oracle” is the result where we know the true active variables and solve on this subset of variables. “glmnet” fits the responses separately. “SRRR-r” indicates the SRRR results with assumed rank *r*.

**Figure 4:**
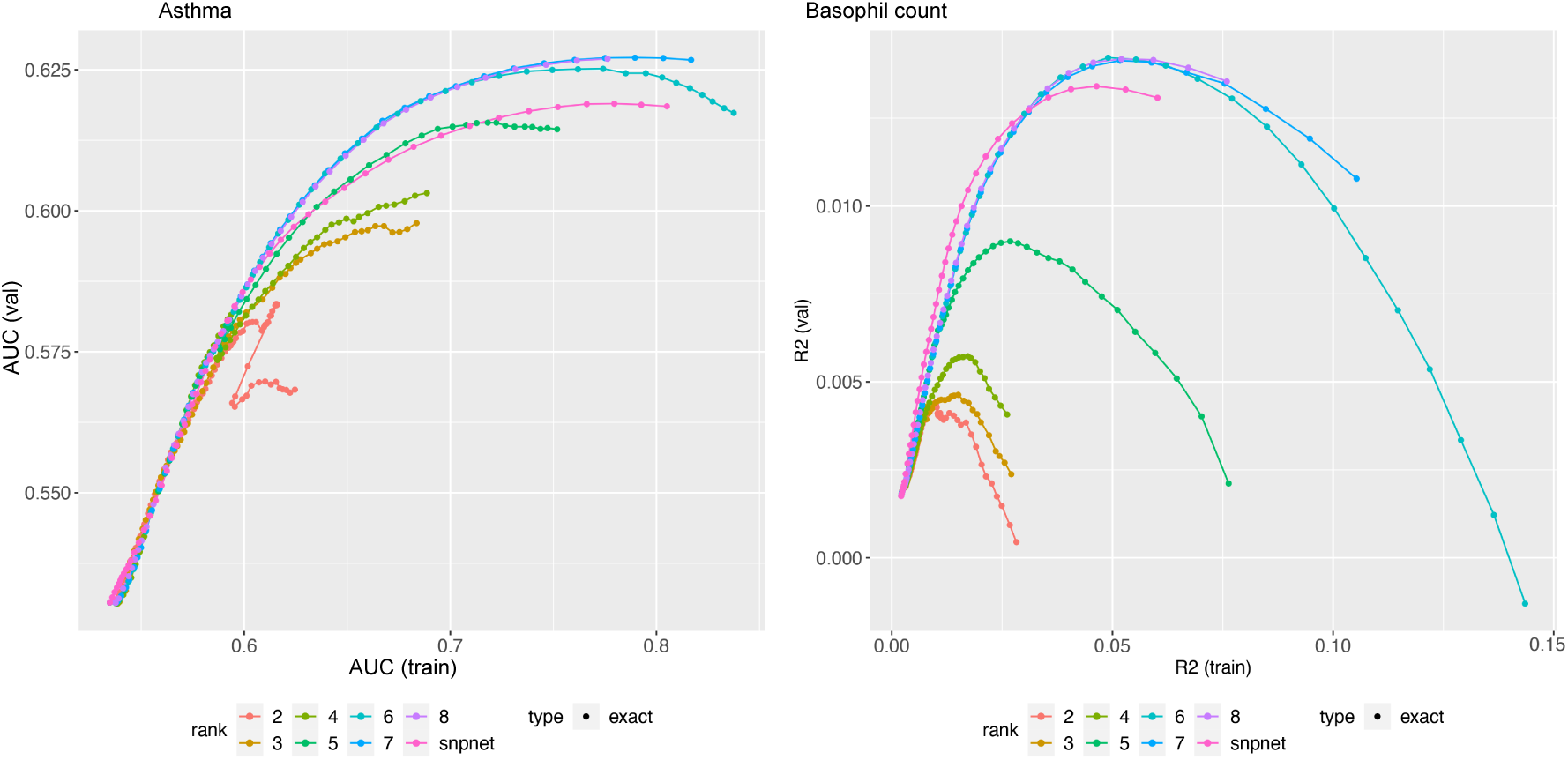
Asthma and Basophil count prediction performance plots. Different colors correspond to lower rank predictive performance across (x-axis) training data set and (y-axis) validation data set for (left) asthma and (right) basophil count.

**Figure 5:**
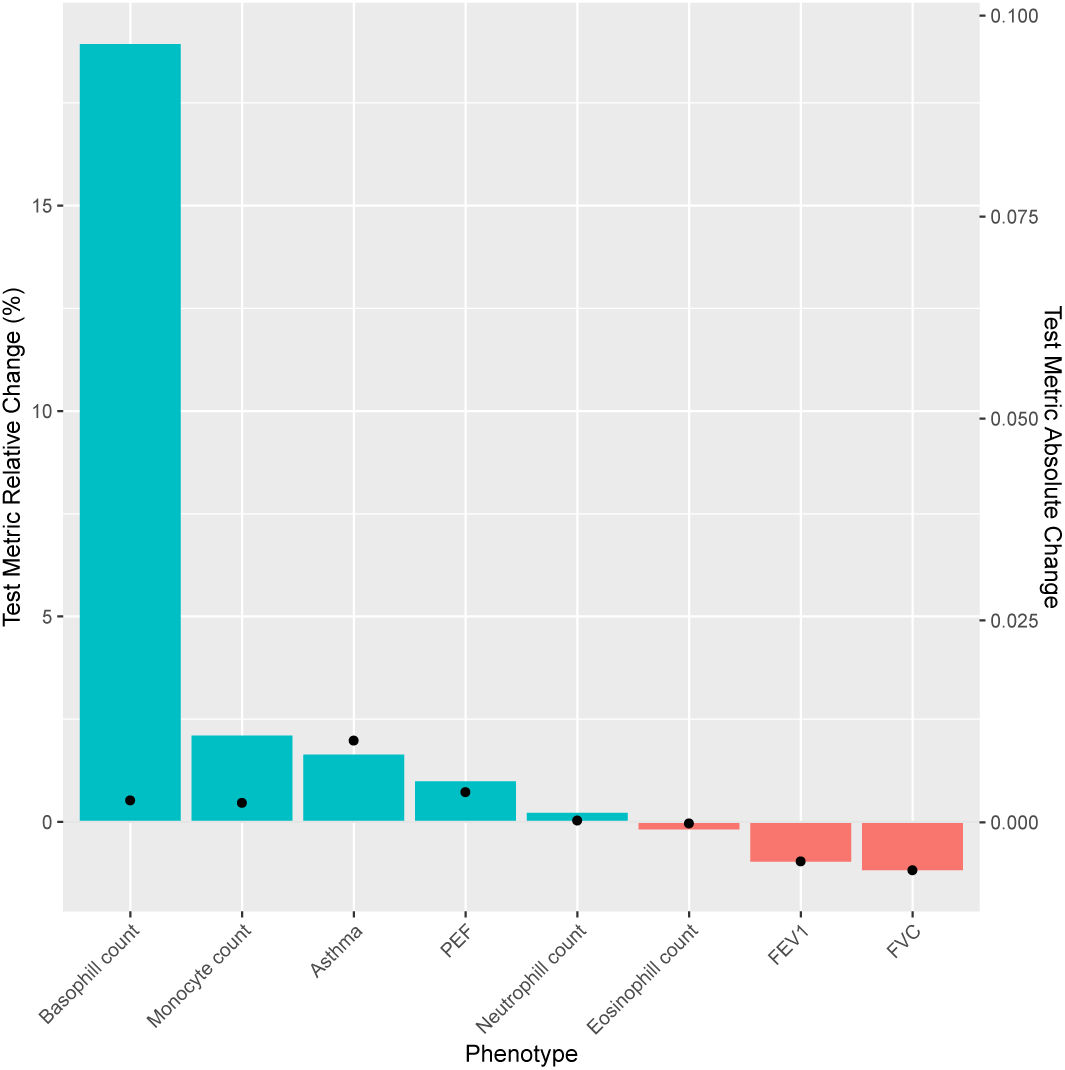
Change in prediction accuracy for multiresponse model compared to single response model. (top) (y-axis 1 bar) *R*^2^ relative change (%) for each phenotype (x-axis) and *R*^2^ absolute change (y-axis 2).

### 6.2 35 Biomarkers

In addition, we used 35 biomarkers from the UK Biobank biomarker panel in Sinnott-Armstrong et al. (2019), and apply SRRR to the dataset. Noticeably, for the liver biomarkers including alanine aminotransferase and albumin, and the urinary biomarkers including Microalbumin in urine and Sodium in urine, we see an improvement in prediction performance for the SRRR application beyond the single-response snpnet models (see Figures 6 and 7).

**Figure 6:**
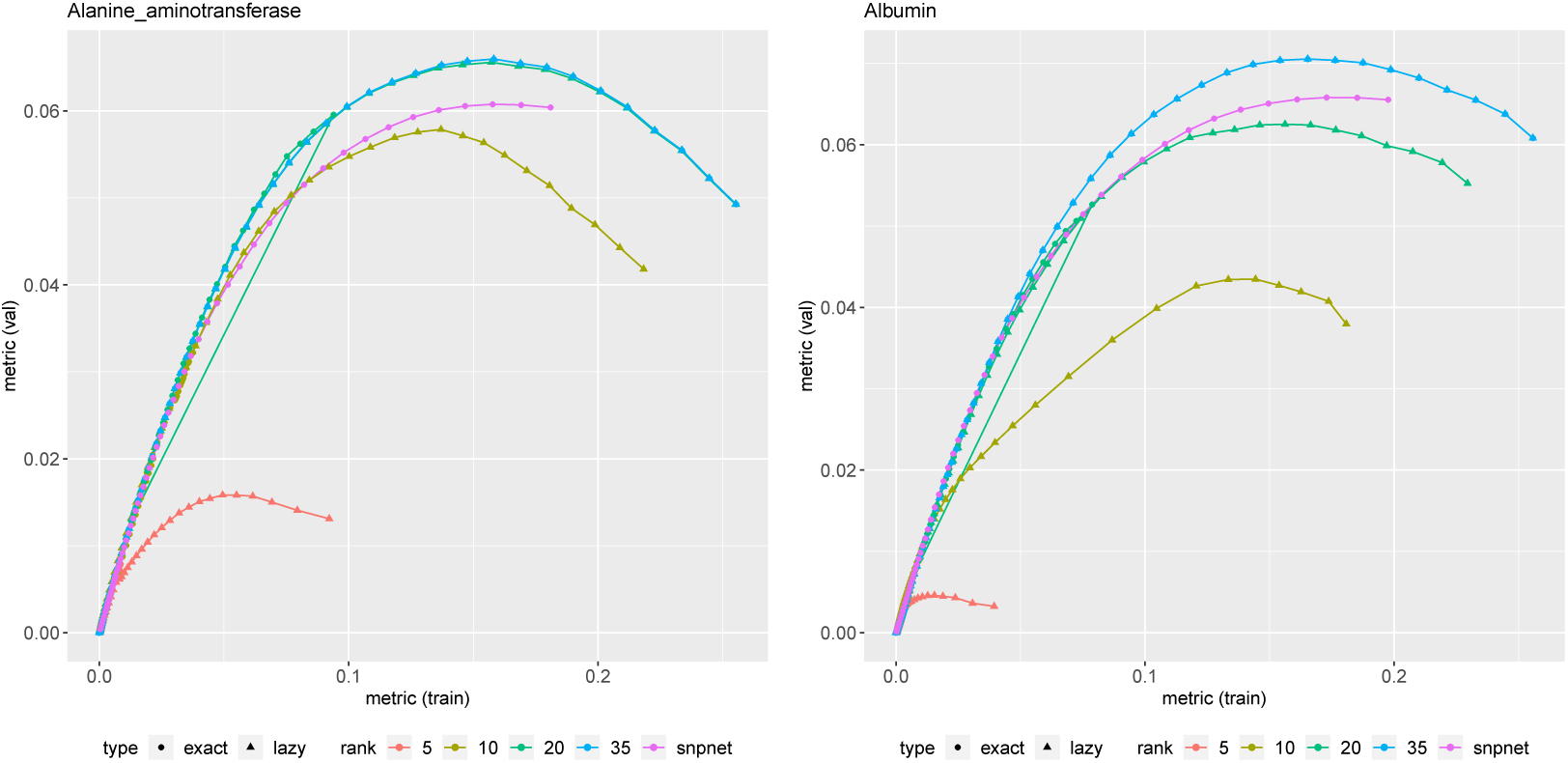
Alanine aminotransferase and albumin prediction performance plots. Different colors correspond to lower rank predictive performance across (x-axis) training data set and (y-axis) validation data set for (left) alanine aminotransferase and (right) albumin. For lower rank representation we applied lazy rank evaluation.

**Figure 7:**
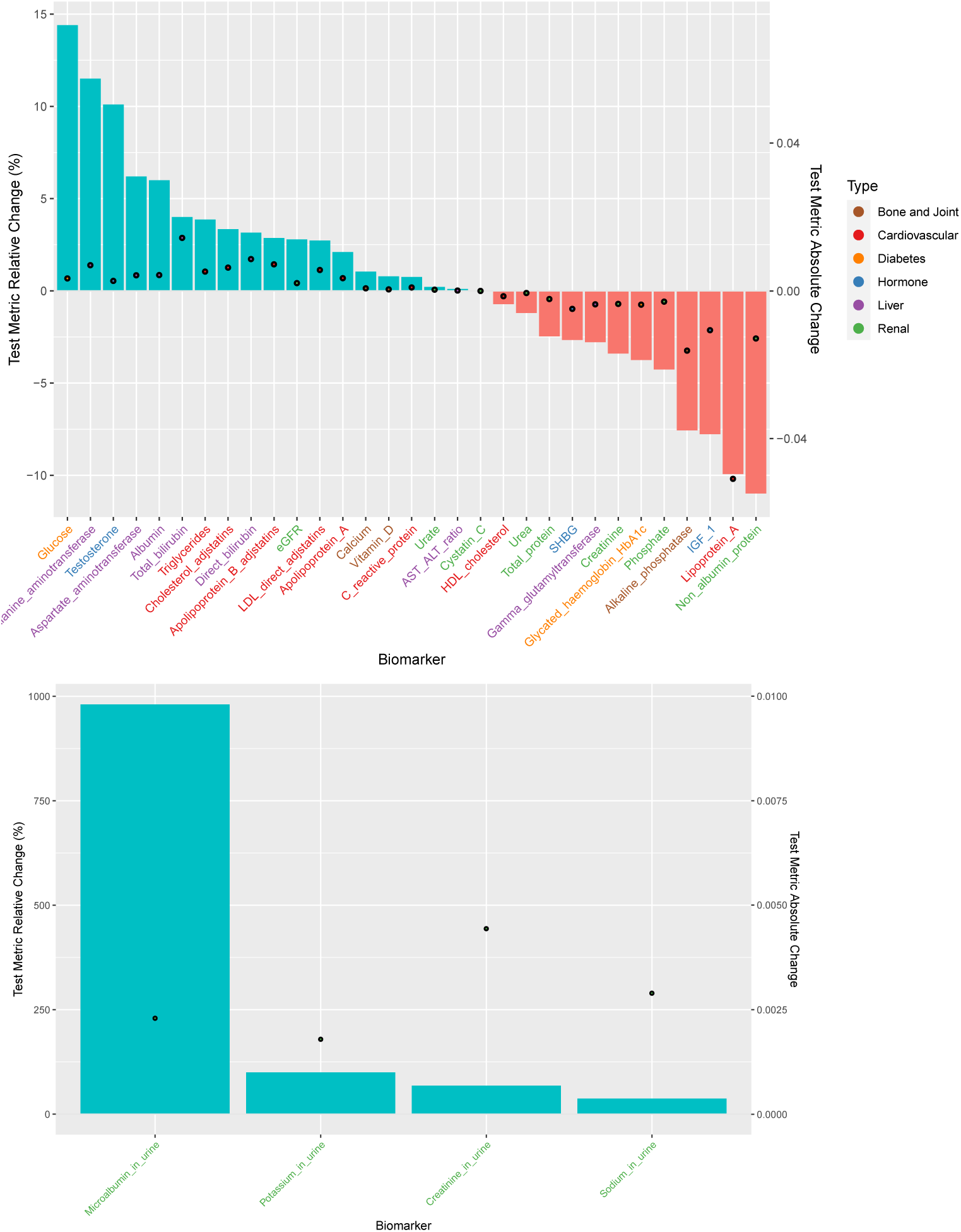
Change in prediction accuracy for multiresponse model compared to single response model. (top) (y-axis 1 bar) *R*^2^ relative change (%) for each biomarker (x-axis) across different biomarker category (color) and *R*^2^ absolute change (y-axis 2). (bottom) Change in predictive accuracy for multiresponse model compared to single response model for urinary biomarkers.

We can represent the lower rank representation as a biplot of the singular value decomposition of the coefficient matrix (Gower et al., 2011; Gabriel, 1971; Tanigawa et al., 2019). Specifically, we display phenotypes projected on phenotype principal components as a scatter plot. We also show variants projected on variant principal components as a separate scatter plot and added phenotype singular vectors as arrows on the plot using sub-axes. In scatter plot with biplot annotation, the inner product of a genetic variant and a phenotype represents the direction and the strength of the projection of the genetic association of the variant-phenotype pair on the displayed latent components. For example, when a variant and a phenotype share the same direction on the annotated scatter plot, that means the projection of the genetic associations of the variant-phenotype pair on the displayed latent components is positive. When a variant-phenotype pair is projected on the same line, but on the opposite direction, the projection of the genetic associations on the shown latent components is negative. When the variant and phenotype vectors are orthogonal or one of the vectors are of zero length, the projection of the genetic associations of the variant-phenotype pair on the displayed latent components is zero. We focused on the top five key SRRR components for AST to ALT ratio (see Figure 8).

**Figure 8:**
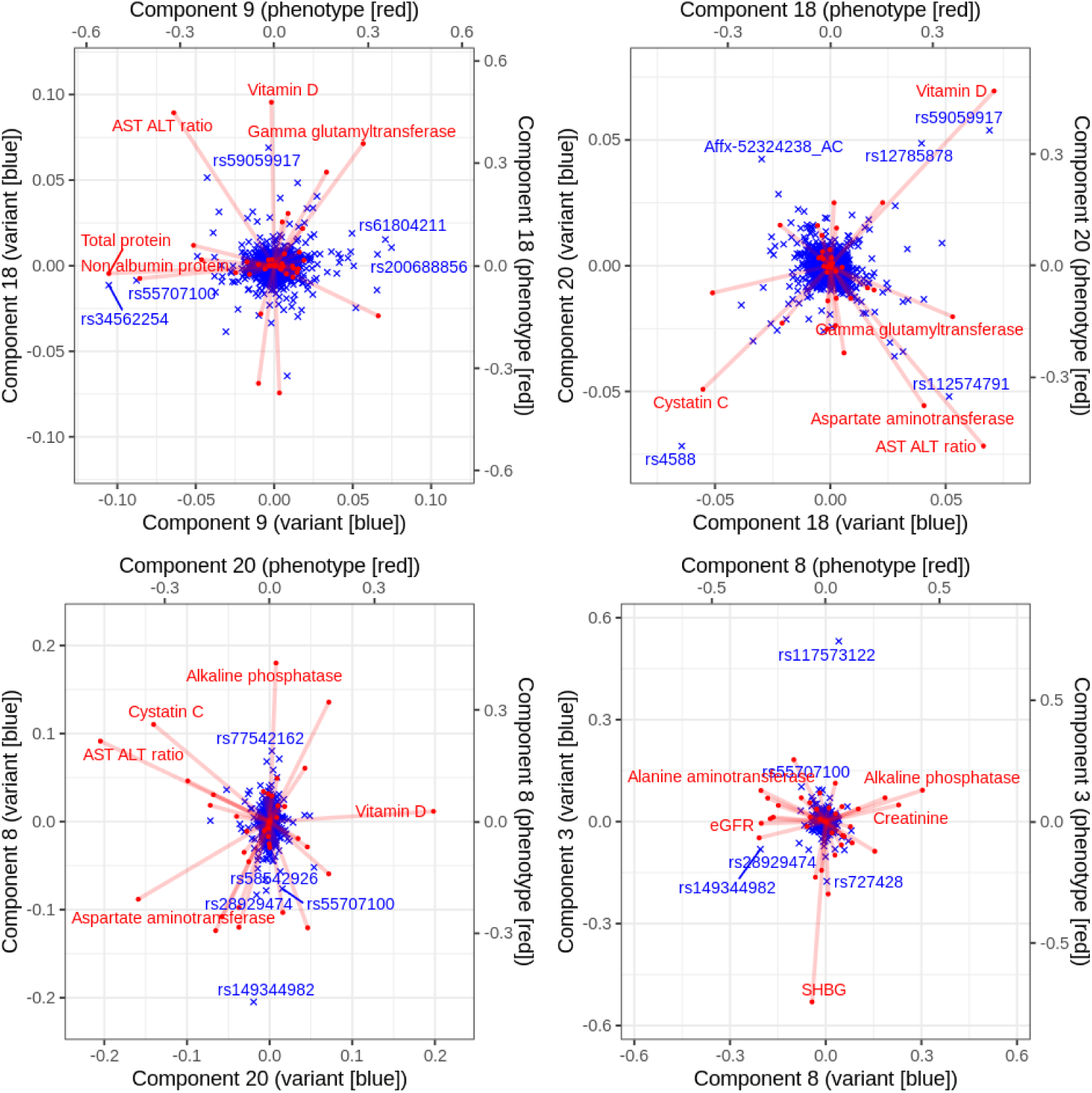
The latent structures of the the top five key SRRR components for AST to ALT ratio. Using trait squared cosine score described in Tanigawa et al. (2019), the top five key SRRR components for AST to ALT ratio (components 9, 18, 20, 8, and 3) are identified from a full-rank SVD of coefficient matrix *C* from SRRR (*C* = *UDV* ^T^) and shown as a series of biplots. In each panel, principal components of genetic variants (rows of *UD*) are shown in blue as scatter plot using the main axis and singular vectors of traits (rows of *V*) are shown in red dots with lines using the secondary axis, for the identified key components. The five traits and variants with the largest distance from the center of origin are annotated with their name.

## 7 Related Work

There are many other methods that were proposed for multivariate regression in high-dimensional settings. Chen, Huang (2012) compares the SRRR with rank-free methods including *L*_2_SVS Similä, Tikka (2007), *L*_*∞*_ SVS (Turlach et al., 2005) that replaces the *ℓ*_2_-norm with *ℓ*_∞_-norm of each row, and RemMap (Peng et al., 2010) that imposes an additional elementwise sparsity of the coefficient matrix. It also compares with the SPLS Chun, Keleş (2010) and points out that the latter does not target directly on prediction of the responses so the performance turns out not as good. Another important category of methods Canonical Correlation Analysis (CCA) (Hotelling, 1936) that tries to constructed uncorrelated components in both the feature space and the response space to maximize their correlation coefficients also falls short in the aspect, even though some connection can be established with the reduced rank regression as seen in Appendix B.

More recently, there is a line of new advances in sparse and low-rank regression problems. For example, Ma, Sun (2014) proposed a subspace assisted regression with row sparsity and studied its near-optimal estimation properties. Ma et al. (2020) furthered this work to a two-way sparsity setting, where nonzero entries are present only on a few rows and columns. Li et al. (2019) proposed an integrative multi-view reduced-rank regression that encourages group-wise low-rank coefficient matrices with a composite nuclear norm. Dubois et al. (2019) developed a fast first-order proximal gradient algorithm on the SRRR objective reparameterized by a single matrix and proves linear local convergence. Luo et al. (2018) proposed a mixed-outcome reduced-rank regression method that deals with different types of responses and also missing data, though it does not aim for high-dimensional settings with variable selection.

In genetics, some approaches proposed to decompose genetic associations from summary level data using LD-pruning along with p-value thresholding for variable selection in an approach referred to as DeGAs (Tanigawa et al., 2019) and MetaPhat (Lin et al., 2019). DeGAs was extended for genetic risk prediction and to “paint” an individual’s risk to a disease based on genetic component loadings in an approach referred to as DeGAs-risk (Aguirre et al., 2019).

## 8 Summary and Discussion

In this paper, we propose a method that takes into account both sparsity in high-dimensional regression problems and low-rank structure when multiple correlated outcomes are present. A screening-based alternating minimization algorithm is designed to deal with large-scale and ultrahigh-dimensional applications, such as the UK Biobank population cohort. We demonstrate the effectiveness of the method on both synthetic and real datasets focusing on asthma and 7 related blood count biomarkers, in addition to the 35 biomarker panel made available by UK Biobank (Sinnott-Armstrong et al., 2019). We anticipate that the approach presented here will generalize to thousands of phenotypes that are currently being measure in UK Biobank, e.g. metabolomics and imaging data that are currently being generated in over 100,000 individuals.

Methodologically, in the UK Biobank experiments, we use continuous approximation to binary outcomes. This is a reasonable assumption but ideally one would like to solve the exact problem based on their respective likelihood. In principle, there is no theoretical challenge in the algorithmic design. We can use Newton’s method and enclose the procedure with an outer loop that conducts quadratic approximation of the objective function. However, the quadratic problem involving both penalty and low-rank constraint can be very messy. We might need some heuristics to find a more convenient approximation. We see this as future work along with extending the SRRR algorithm to other families including time-to-event multiple responses that can be used for survival analysis. Furthermore, for an individual we can project a variant and phenotype loading across the reduced rank to their risk to arrive at a similar analysis of outlier individuals with unusual painting of genetic risk and to quantify the overall contribution of a component which may aid in disease risk interpretation. Overall, we see the method and algorithms presented here as an important toolkit to the prediction problem in human genetics.

## Acknowledgement

This research has been conducted using the UK Biobank Resource under Application Number 24983, “Generating effective therapeutic hypotheses from genomic and hospital linkage data” (http://www.ukbiobank.ac.uk/wp-content/uploads/2017/06/24983-Dr-Manuel-Rivas.pdf). Based on the information provided in Protocol 44532 the Stanford IRB has determined that the research does not involve human subjects as defined in 45 CFR 46.102(f) or 21 CFR 50.3(g). All participants of UK Biobank provided written informed consent (more information is available at https://www.ukbiobank.ac.uk/2018/02/gdpr/). We thank all the participants in the UK Biobank. M.A.R. is supported by Stanford University and a National Institutes of Health (NIH) Center for Multi- and Trans-ethnic Mapping of Mendelian and Complex Diseases grant (5U01 HG009080). Y.T. is supported by a Funai Overseas Scholarship from the Funai Foundation for Information Technology and the Stanford University School of Medicine. Research reported in this publication was supported by the National Human Genome Research Institute of the NIH under Award Number R01HG010140 (M.A.R.). The content is solely the responsibility of the authors and does not necessarily represent the official views of the NIH. R.T. was partially supported by NIH grant 5R01 EB001988-16 and NSF grant 19 DMS1208164. T.H. was partially supported by grant DMS-1407548 from the National Science Foundation, and grant 5R01 EB 001988-21 from the National Institutes of Health.

## A Additional Proofs

### A.1 Proof of Lemma 1

This is intuitively the same as one without the rank constraint because when the coefficients just start to become nonzero, the coefficient matrix is low-rank in its nature. Therefore, for the purpose of finding the maximum meaningful *λ*, we can ignore the rank constraint unless *r* = 0. Without the constraint, it follows from the KKT condition that having all coefficients to be zero is equivalent to setting

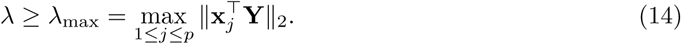

Therefore, the maximum *λ* that accommodates a nontrivial solution is 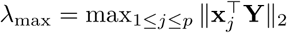.

### A.2 Proof of Lemma 2

We plug in the SVD of **Z** and have 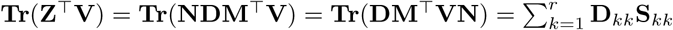 where **S** = **M**^T^**VN** and the last equality is due to the fact that **D** is a diagonal matrix. Notice that by the skinny SVD, **SS**^T^ = **M**^T^**V**^T^**NN**^T^**VM** = **I**. We thus know **S** is an orthogonal matrix and the magnitude of its diagonal elements cannot exceed 1. Since **D**_*kk*_ are all non-negative. To maximize 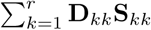, we let **S**_*kk*_ = 1 for all 1 ≤ *k* ≤ *r*. This is equivalent to setting **S** = **M**^T^**VN** = **I**. Therefore, one solution is given by **V** = **MN**^T^. The maximum value of the objective is thus 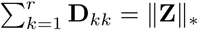, the nuclear norm of **Z**.

### A.3 Proof of Theorem 2

We notice that in Problem (3) we can solve explicitly for **V** and plug back into the objective function. It yields the objective function (after dropping the constant term 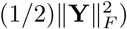:

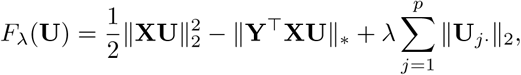

We let 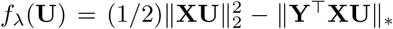 without the penalty term so that 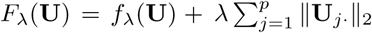. Define a local smooth approximation of *F*_*λ*_ as

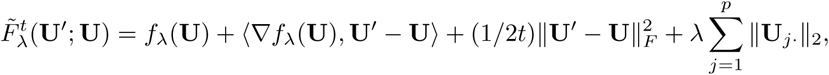

and 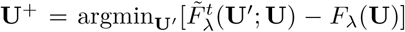. Dubois et al. (2019) showed that if *t* is small enough such that 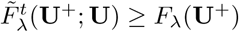, we have

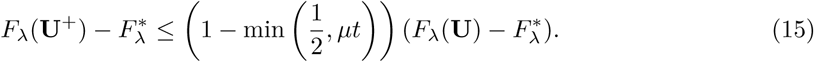

Consider the iterates (**U**^*k*^, **V**^*k*^)_*k*≥1_ in the alternating minimization algorithm. Notice that ∇*f*_*λ*_(**U**^*k*^) = **X**^T^**XU**^*k*^ − **X**^T^**YV**^*k*^. We have

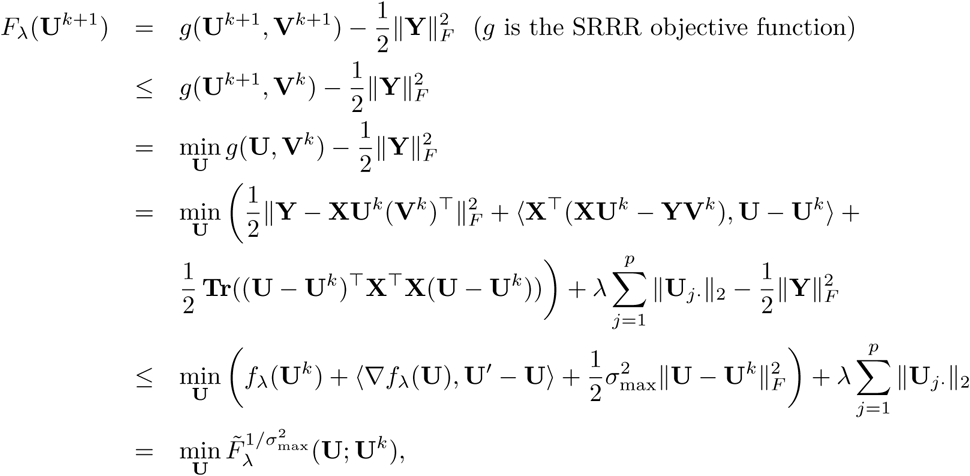

where the fourth line is the quadratic expansion of *g*(**U, V**^*k*^) at **U**^*k*^, the second to last is by the fact that 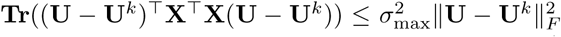, and the last equality is by the definition of 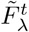 function. Therefore, if we let 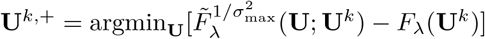, we have

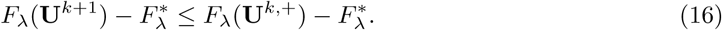

We need to show that **U**^*k*,+^ satisfies the condition 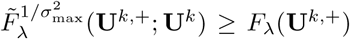. To see this, notice that in fact for any **U**,

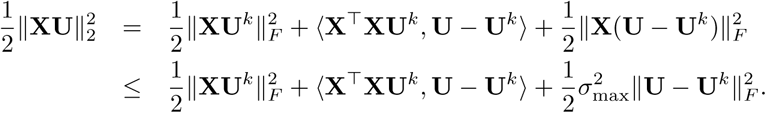

Since **X**^T^**YV**^*k*^ is a subgradient of ‖**Y**^T^**XU** ‖ _*\**_ at **U**^*k*^, we have

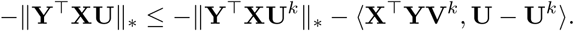

Adding the two inequalities up, and we have 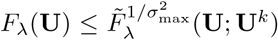 for all **U**. In particular, it holds for **U**^*k*+^. Therefore, by (15) and (16), we have

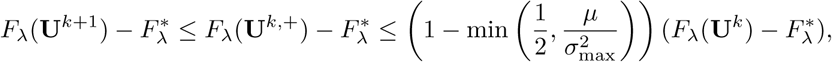

and the convergence is linear.

## B Connection with CCA

Canonical Correlation Analysis (CCA) has an internal connection with Reduced-Rank Regression (RRR). In particular, it can be shown that the low-rank components constructed on the **X** space turn out to be the same by a relaxed CCA and a generalized RRR. CCA finds linear combinations **XU** ∈ ℝ^*n*×*r*^ of variables in **X** ∈ ℝ^*n*×*p*^ and linear combinations **YV** ∈ ℝ^*n*×*r*^ of variables in **Y** ∈ ℝ^*n*×*q*^ that attain the maximum correlation. We assume both **X** and **Y** have been centered. CCA solves the following optimization problem:

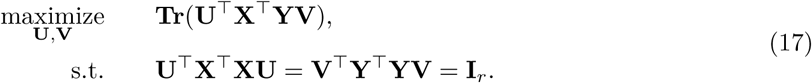

In particular, in the one dimensional case, this reduces to the problem of maximizing our familiar correlation coefficient. An equivalent representation to (17) can be written as

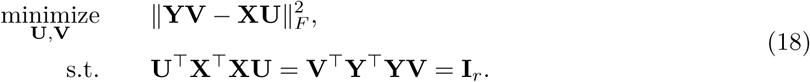

The solution to the problem is 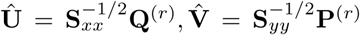where **P**^(*r*)^ and **Q**^(*r*)^ are the *r* leading left and right singular vectors of matrix 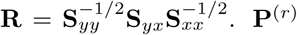 is also the *r* leading eigenvectors of 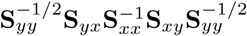. A relaxed form of CCA problem ignoring the **U**-constraint solves

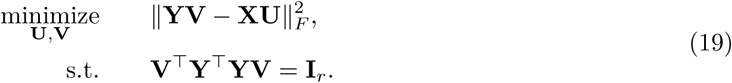

The solution is 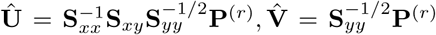, where **P**^(*r*)^ is the *r* leading eigenvectors of 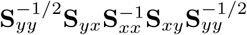. Therefore, the solution for **V** remains unchanged, though **U** is different due to the constraint.

On the other hand, in the (generalized) reduced rank regression, given a given positive-definite matrix Γ, the problem becomes

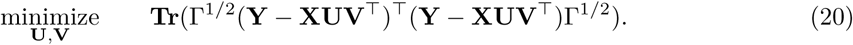

This can be derived, for example, as an maximum likelihood estimator under the Gaussian assumption with known covariance Γ^−1^. One solution (Velu, Reinsel, 2013) is given by

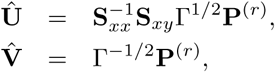

where **P**^(*r*)^ is the leading eigenvectors of 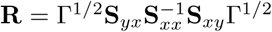. We see that the solution when 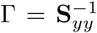 is closely related to the relaxed CCA solution. **U** is the same while **V** is the so-called reflexive inverse of **V** there.

## C Additional Experiments

We conduct some experiments to gain more insight into the method and compare with other methods. We generate the **X** ∈ ℝ^*n*×*p*^ with independent samples from some multivariate Gaussian 𝒩(0, Σ_*X*_). For the first several cases, we generate the response from the true, most favorable model **Y** = **XUV**^T^ + **E**, where each entry in the support of **U** ∈ ℝ^*p*×*r*^ (sparsity *k*) is independently drawn from a standard Gaussian distribution, and **V** ∈ ℝ^*q*×*r*^ takes the left singular matrix of a Gaussian ensemble. Hence **B** = **UV**^T^ is the true coefficient matrix. The noise matrix is generated from 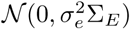, where 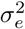 is chosen such that the signal-to-noise ratio

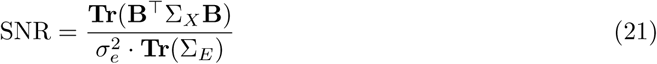

is set to a given level. The performance is evaluated by the test *R*^2^, defined as follows:

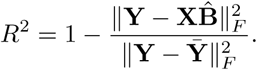

We consider several sets of experiments.

1. **Scenario 1-9** Small experiments: (*n, p, k*) = (200, 100, 20), (200, 500, 20), (200, 500, 50), *q* = 20, *r* = 3. The *X* has independent design, and the noise across different responses are all independent, i.e. Σ_*X*_ = **I**_*p*_, Σ_*E*_ = **I**_*q*_. Target SNR = 0.5, 1, 3. The results are evaluated on test sets of size 5000.
2. **Scenario 10-18** Same as Scenario 1-9. The true coefficient matrix is no longer exact low rank. It is perturbed by Gaussian noise with mean 0 and standard deviation 0.5.
3. **Scenario 19-27** Same as Scenario 1-9, except that the predictors are correlated. In particular,

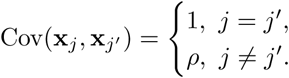 We let *ρ* = 0.5 in this set of simulation.
4. **Scenario 28-36** Same as Scenario 10-18, except that the predictors are correlated as in Scenario 19-27.

From the simulations, we find that underestimating the rank can degrade the performance instantly. Overestimating the rank will give one a variance penalty, but it seems to be rather robust compared with the other direction.

**Scenario 1-9** Small experiments: (*n, p, k*) = (200, 100, 20), (200, 500, 20), (200, 500, 50), *q* = 20, *r* = 3. The **X** has independent design, and the noise across different responses are all independent, i.e. Σ_*X*_ = **I**_*p*_, Σ_*E*_ = **I**_*q*_. Target SNR = 0.5, 1, 3. The results are evaluated on test sets of size 5000.

**Figure C.1:**
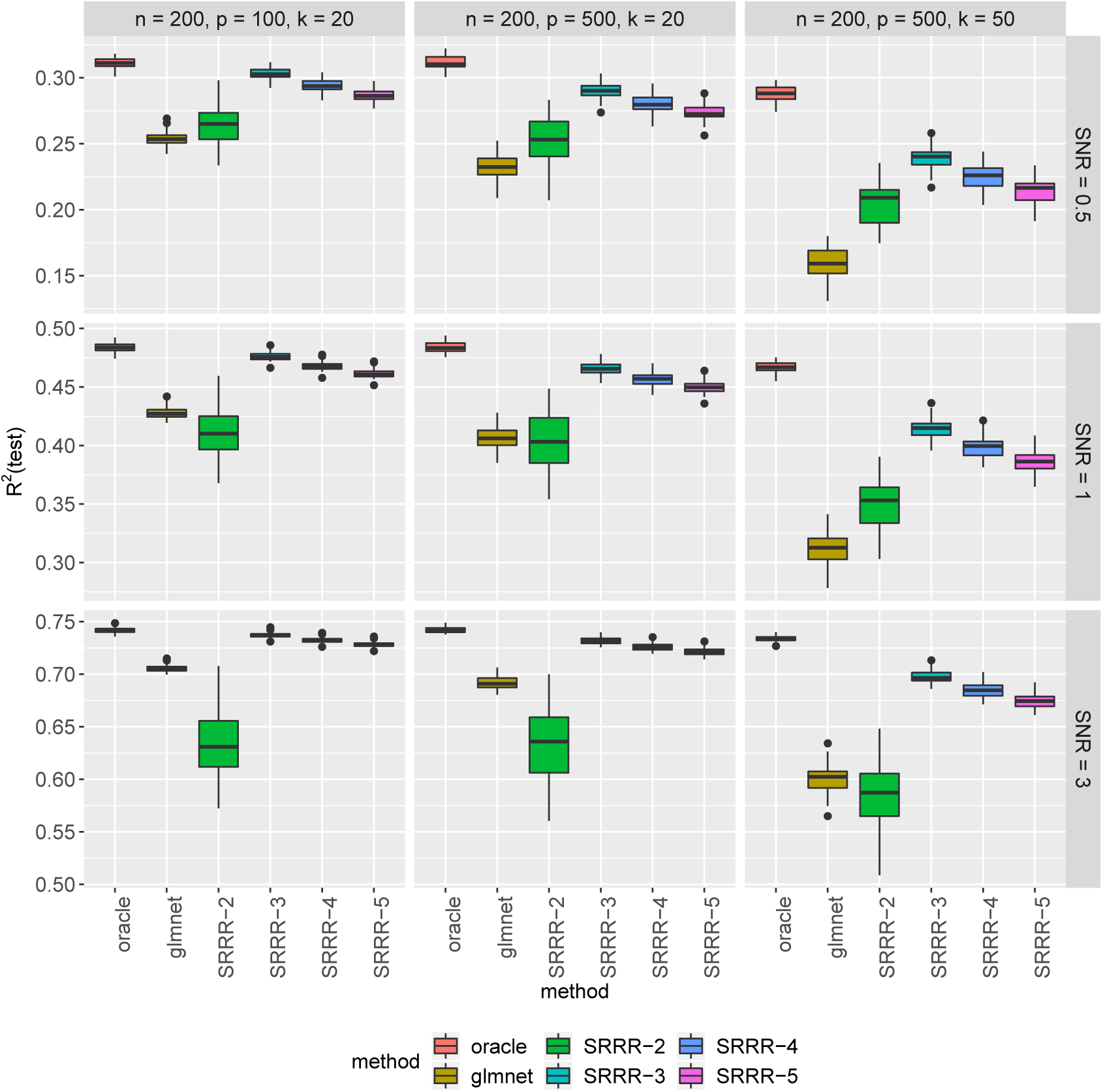
Scenario 1-9. *R*^2^ each run is evaluated on a test set of size 5000.

**Scenario 10-18** Same as Scenario 1-9. The true coefficient matrix is no longer exact low rank. It is perturbed by Gaussian noise with mean 0 and standard deviation 0.5.

**Figure C.2:**
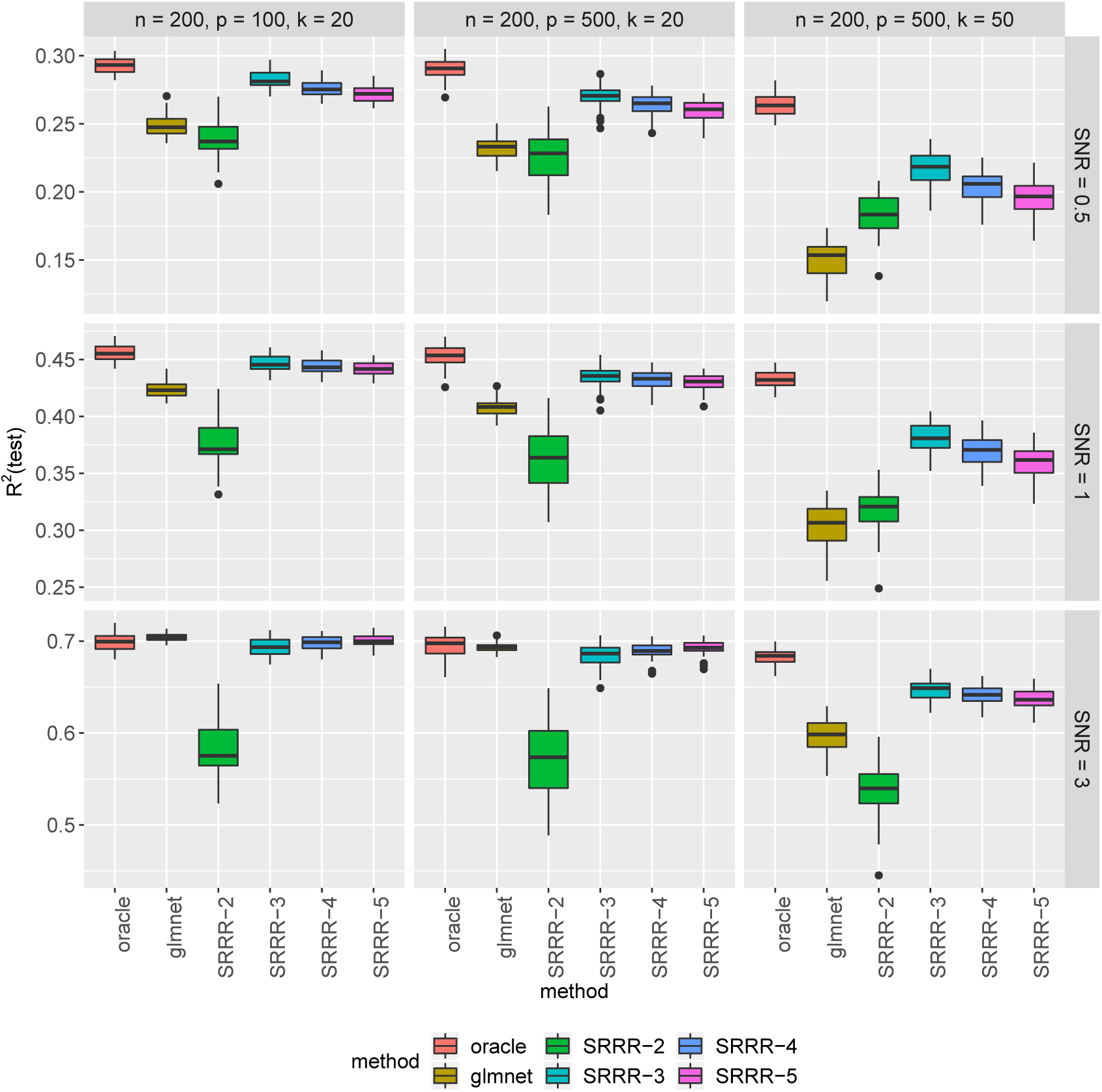
Scenario 10-18. *R*^2^ each run is evaluated on a test set of size 5000. The oracle here does not take into account the noise in true coefficient matrix, and do reduced rank regression on the true support and the true rank.

**Scenario 19-27** Same as Scenario 1-9, except that the predictors are correlated. In particular,

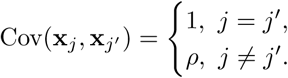

We let *ρ* = 0.5 in this set of simulation.

**Figure C.3:**
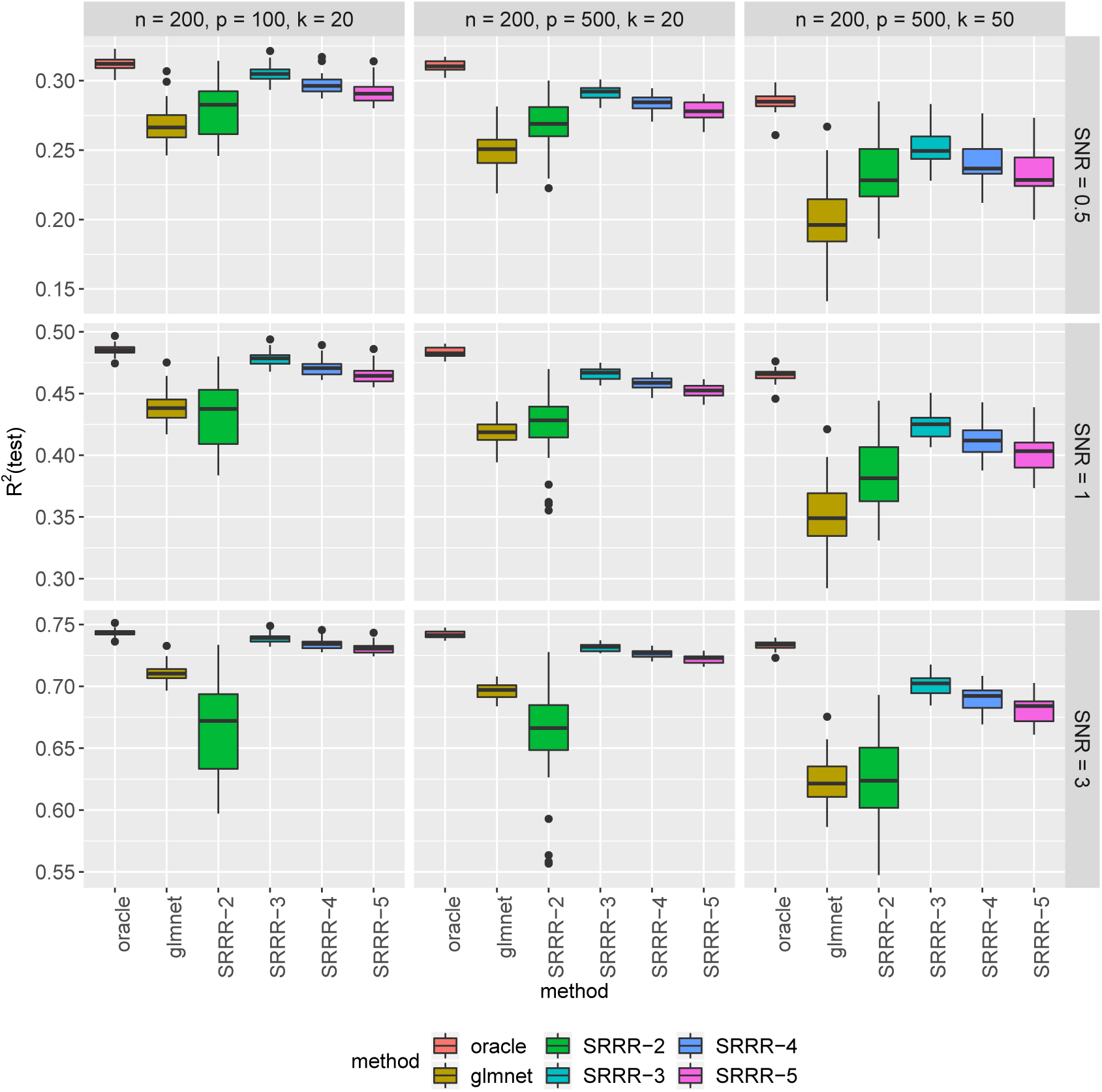
Scenario 19-27. *R*^2^ each run is evaluated on a test set of size 5000.

**Scenario 28-36** Same as Scenario 10-18, except that the predictors are correlated as in Scenario 19-27.

**Figure C.4:**
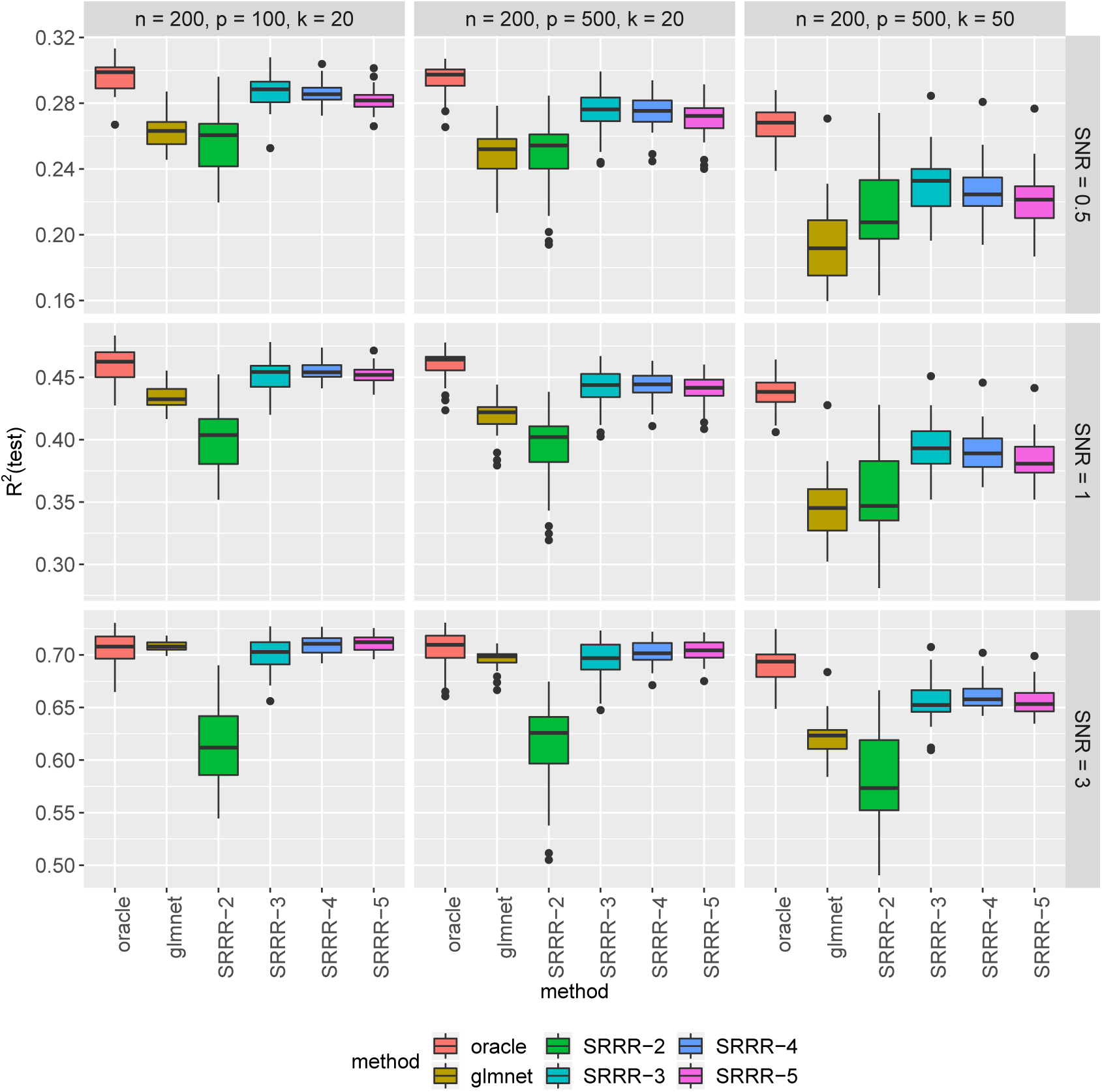
Scenario 28-36. *R*^2^ each run is evaluated on a test set of size 5000.

## D Additional Information on the Methods

### D.1 Compliance with ethical regulations and informed consent

This research has been conducted using the UK Biobank Resource under Application Number 24983, “Generating effective therapeutic hypotheses from genomic and hospital linkage data” (http://www.ukbiobank.ac.uk/wp-content/uploads/2017/06/24983-Dr-Manuel-Rivas.pdf). Based on the information provided in Protocol 44532 the Stanford IRB has determined that the research does not involve human subjects as defined in 45 CFR 46.102(f) or 21 CFR 50.3(g). All participants of UK Biobank provided written informed consent (more information is available at https://www.ukbiobank.ac.uk/2018/02/gdpr/).

### D.2 Population stratification in UK Biobank

We used genotype data from the UK Biobank dataset release version 2 and the hg19 human genome reference for all analyses in the study. To minimize the variabilities due to population structure in our dataset, we restricted our analyses to include 337,151 White British individuals (Figure D.1) based on the following five criteria (DeBoever et al., 2018; Tanigawa et al., 2019) reported by the UK Biobank in the file “ukb_sqc_v2.txt”:

1. self-reported white British ancestry (“in_white_British_ancestry_subset” column)
2. used to compute principal components (“used_in_pca_calculation” column)
3. not marked as outliers for heterozygosity and missing rates (“het missing outliers” column)
4. do not show putative sex chromosome aneuploidy (“putative_sex_chromosome_aneuploidy” column)
5. have at most 10 putative third-degree relatives (“excess_relatives” column).

### D.3 Variant annotation and quality control

We prepared a genotype dataset by combining the directly-genotype variants, copy number variants (CNVs) and HLA allelotype datasets.

We annotated the directly-genotyped variants using the VEP LOFTEE plugin (https://github.com/konradjk/loftee) and variant quality control by comparing allele frequencies in the UK Biobank and gnomAD (gnomad.exomes.r2.0.1.sites.vcf.gz) as previously described28. We focused on variants outside of the major histocompatibility complex (MHC) region (chr6:25477797-36448354) as previously described. We focused on the variants according to the following criteria:

- Missigness of the variant is less than 1%, considering that two genotyping arrays (the UK BiLEVE array and the UK Biobank array) which covers a slightly different set of variants.
- Minor-allele frequency is greater than 0.01%, given the recent reports casting questions on the reliability of ultra low-frequency variants.
- The variant is in the LD-pruned set
- Hardy-Weinberg disequilibrium test p-value is less than 1.0 × 10^−7^
- Manual cluster plot inspection. We investigated the cluster plots for subset of variants and removed 11 variants that have unreliable genotype calls.
- Passed the comparison of minor allele frequency with gnomAD dataset as described before

CNVs were called by applying PennCNV v1.0.4 on raw signal intensity data from each array within each genotyping batch as previously described. We applied a filter on minor-allele frequency (MAF > 0.01%), which resulted in 8,274 non-rare (MAF > 0.01%) CNVs.

The HLA data from the UK Biobank contains all HLA loci (one line per person) in a specific order (A, B, C, DRB5, DRB4, DRB3, DRB1, DQB1, DQA1, DPB1, DPA1). We downloaded these values, which were imputed via the HLA:IMP*2 program (Resource 182); the UK Biobank reports one value per imputed allele, and only the best-guess alleles are reported. Out of the 362 alleles reported in UKB, we used 175 alleles that were present in >0.1% of the population surveyed.

**Figure D.1:**
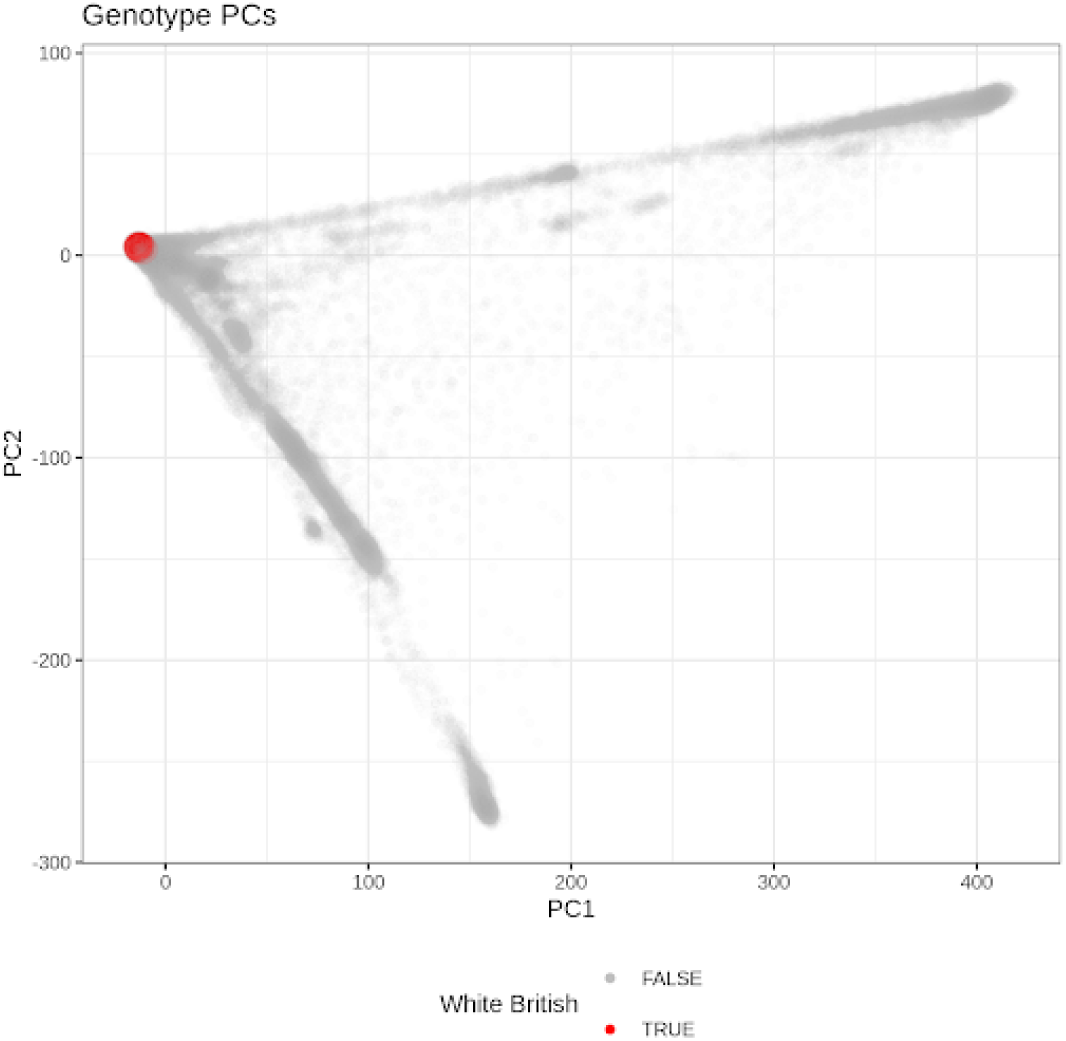
The identification of unrelated White British individuals in UK Biobank. The first two genotype principal components (PCs) are shown on the x- and y-axis and the identified unrelated White British individuals (Methods) are shown in red.

## Footnotes

1 We use 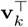 to represent the *k*th row of **V** for convenience.

## Notes

### Competing Interest Statement

The authors have declared no competing interest.

